# Inference of elevated mutation rates and variant effects using 700k exomes

**DOI:** 10.64898/2026.06.09.730991

**Authors:** Prathitha Kar, Mikhail A. Moldovan, Jeremy Guez, Sumaiya Nazeen, Julia K. Goodrich, Trisha Karani, Kaitlin E. Samocha, Konrad J. Karczewski, Evan Koch, Vladimir Seplyarskiy, Shamil R. Sunyaev

## Abstract

Genomic sequencing is now widely accessible for genetic diagnostics and is emerging as a component of newborn screening. This technological development generates the need to characterize incoming mutations, create comprehensive datasets of genes causing rare Mendelian disorders, and identify pathogenic variants. Large-scale exome sequencing datasets such as Genome Aggregation Database (gnomAD) have been assembled to help address these challenges. The recent release of gnomAD (v4; n = 730,947) uncovers millions of rare coding variants, many of which have arisen more than once by independent recurrent mutations in the rapidly growing recent human population. Here, we use newly developed theoretical understanding of sampling properties of rare variants to estimate key population genetics parameters of practical importance to human genetics such as demography history, mutation rate, and selection. Solely relying on population data, our method Population Inferred Estimates of Selection (PIES) identifies novel genes with loss-of-function mutational hotspots likely due to selection in spermatogonia. PIES efficiently estimates selection coefficients for heterozygous loss-of-function variants. Combining population genetics inference with variant effect predictors, PIES predicts pathogenic missense mutations and improves variant prioritization for genetic diagnostics and newborn screening.

## Introduction

On average, a human newborn carries 70 *de novo* single-nucleotide mutations [1, 2]. Of special importance are mutations in the protein coding regions of the genome; up to 1 in 300 newborns develop a monogenic disease caused by a *de novo* mutation in a protein coding gene [3, 4]. Mutational influx also generates a myriad of segregating allelic variants in the human population, most of which are very rare [5]. The ongoing efforts in the analysis and interpretation of rare variation are key to improving disease gene discovery, diagnostics, and more generally, our understanding of the impact of genetic variation on biological function. Despite the progress, knowledge gaps in the functional, evolutionary, and clinical characterization of newly arising mutations and segregating rare variants persist.

Most rare variants detectable in large sequencing datasets arose relatively recently in the rapidly growing human population. The large size of the population produced a large supply of mutations. As a result, many rare variants arose by mutation more than once (the phenomenon known as mutational recurrence) [6–8]. The rate of mutational recurrence is expected to be higher for sites with elevated *de novo* mutation rates [9]. Multiple mutations at a site increase allele frequencies, introducing coupling between mutation rate and frequency for rare variation: sites with higher *de novo* mutation rates harbor variants at higher sample counts in large population cohorts than sites with lower mutation rates [5, 10, 11].

Recent progress in the theoretical understanding of sampling properties of rare variants, together with improved estimates of *de novo* mutation rates, has enabled prediction of the frequency distribution of rare variation in humans with high precision [7, 9, 12–14]. Here, we combine these advances using a Bayesian framework called PIES (**P**opulation **I**nferred **E**stimates of **S**election) to estimate gene and variant-specific population genetics parameters of key interest to human genetics (Figure 1). We apply PIES to allele frequencies extracted from 730,947 exomes and made publicly available in the Genome Aggregation Database (gnomAD) v4 dataset [5]. PIES incorporates basepair-resolution mutation rate estimates from the Roulette model and explicitly accounts for uncertainty in its predictions [12].

**Figure 1:**
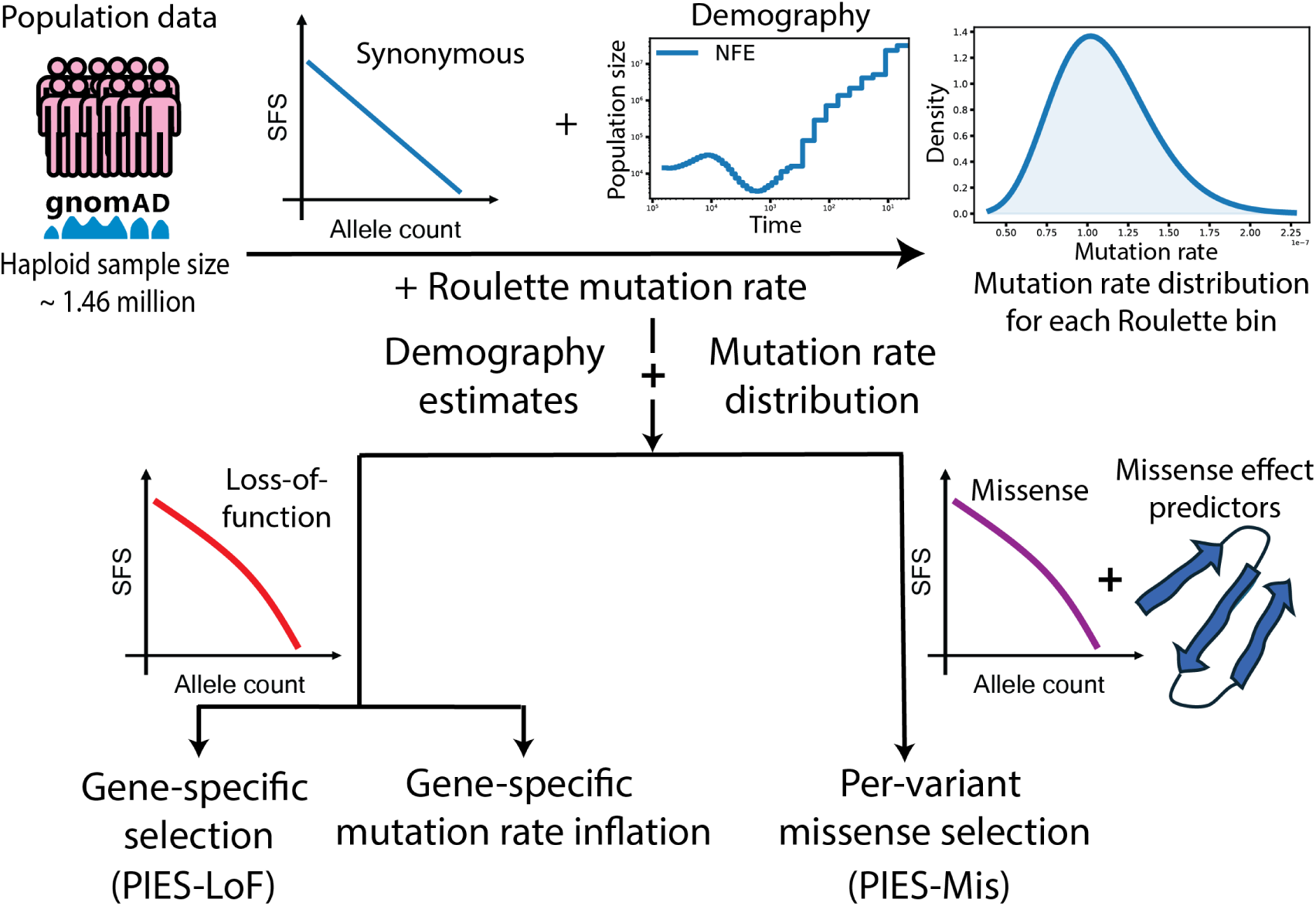
Overview of the Population Inferred Estimates of Selection (PIES) framework. PIES analyzes large-scale population sequencing data from gnomAD v4 to infer key population genetic parameters, including mutation rates, demographic history, and selection. Mutation rates and their uncertainty are estimated using the site frequency spectrum (SFS) of synonymous variants, together with an ancestry-specific demography (such as for Non-Finnish Europeans) and the Roulette mutation rate model. These estimates are then fixed as global parameters in downstream analyses. Using this calibrated model, PIES infers gene-level selection acting against loss-of-function (LoF) variants from the LoF SFS (PIES-LoF). Recent studies have found genes whose LoFs have a higher mutation rate than expected under baseline mutation rate models due to clonal expansions in spermatogonia. We use PIES to find such genes using population data. Finally, PIES extends to missense variation (PIES-Mis) by integrating missense SFS with existing variant effect predictors.

The first quantity we estimate is the strength of selection against heterozygous loss-of-function (LoF) variants. Measures of selective constraint [10, 11] and estimates of heterozygous selection coefficients [8, 15–18] of LoF variants have found applications in discovery of autosomal dominant disease genes [19–21]. Highly selectively constrained genes are predominantly involved in critical biological functions and enriched in heritability of complex phenotypes [22, 23]. Several methods have been developed to estimate selection coefficients or related measures of selective constraint against heterozygous LoF variants [8, 10, 11, 15–18]. Here, we provide new estimates of gene-specific LoF selection coefficients and show their utility in prioritizing haploinsufficient genes causing monogenic diseases. These improved estimates also show the scalability of PIES to large datasets such as gnomAD v4.

An interesting consequence of LoF mutations is their role in clonal expansion in tissues. Recent studies using trio sequencing and direct sperm sequencing found that LoF mutations causing clonal expansions in spermatogonia (CES) had the appearance of elevated mutation rates [24, 25]. CES were originally discovered for gain-of-function mutations leading to severe early onset neurodevelopmental conditions [26–30]. Recently, CES has also been documented for LoF mutations [24, 25] including those that do not cause obvious monogenic dominant developmental diseases with high penetrance [24]. In order to identify additional LoF mutation hotspots genes, we extended our approach to detect elevated effective mutation rates using population data alone. Our analysis identified six known and predicted three new genes hypermutable for LoF variants. These novel genes have biological properties consistent with their role in spermatogonia.

Arguably, the most practically important problem in the interpretation of rare coding variation is the prediction of deleterious missense variants [31]. Missense changes are the most abundant class of protein-altering variants. They have a wide range of possible effects ranging from completely benign to severe. The enormous challenges in predicting the deleteriousness of variants hinder the characterization of variants of uncertain significance (VUS) [32]. Two advances have helped to address these challenges - first, tabulation of population genomic datasets provides information about the selection acting against variants. Secondly, over the span of two decades, the field developed many computational methods for predicting damaging missense variants (such as deep learning models - AlphaMissense, ESM-1b, EVE, popEVE, PrimateAI-3D, MisFit [33–38] or more traditional methods - Polyphen, SIFT, VEST, CADD, REVEL, VARITY [39–45]). We address the challenge of systematically combining these two complementary sources of information. PIES-Mis estimates selection coefficients of individual missense variants by integrating existing variant effect predictors with our population genetics framework. For each gene, we fit just three parameters, enabling inference without requiring expensive training. This allows the model to be continuously recomputed to incorporate new sequencing datasets and variant effect predictors. Learning from 730,947 exomes, it achieves a higher accuracy than available methods across multiple benchmarks including predicting variants involved in neurodevelopmental disease.

## Results

### PIES model

Site frequency spectrum (SFS) is simultaneously influenced by mutation rate, demographic history, and selection. The identification of mutation hotspots from population variation data requires the proper context of demography and mutation rate. Similarly, accurate inference of selection requires reliable estimates of mutation rates and demography. We develop PIES to estimate these population genetics parameters from the SFS of rare variants (Figure 1). PIES is designed for the analysis of large-scale datasets, such as gnomAD v4, where recurrent mutations are common, affecting the distribution of rare variants [6–8]. It uses recent methodological development - DR EVIL - to calculate the likelihood of observed SFS [14].

PIES conventionally employs a parametrized model of demographic history [46, 47] capturing changes of effective population size (*N_e_*) over time. For given ancestry, parameters of population history are treated as global rather than gene specific (all variants in all genes follow the same history). Demographic history estimates for populations of different ancestry are shown in Supplementary Figure S1 and Supplementary Table S1 (see Methods for details).

As noted above, presence of recurrent mutation couples mutation rate with the shape of SFS (Figures 2A,B). PIES is supplied with the mutation rate model Roulette [12]. PIES rescales Roulette estimates to per generation per site rates using the SFS at synonymous sites, stratified by mutation rate (Supplementary Figure S2 and Supplementary Table S2, see Methods). Critically, PIES evaluates uncertainty of mutation rate estimates using the same approach as described in ref [14] (Supplementary Figure S2 and Supplementary Table S2). We found that inclusion of the uncertainty in the model substantially improves goodness-of-fit (Figure 2A,B and Supplementary Figure S2) and estimation of the proportion of segregating sites (Figure 2C). Therefore, PIES incorporates mutation rate uncertainty in all downstream inferences.

**Figure 2:**
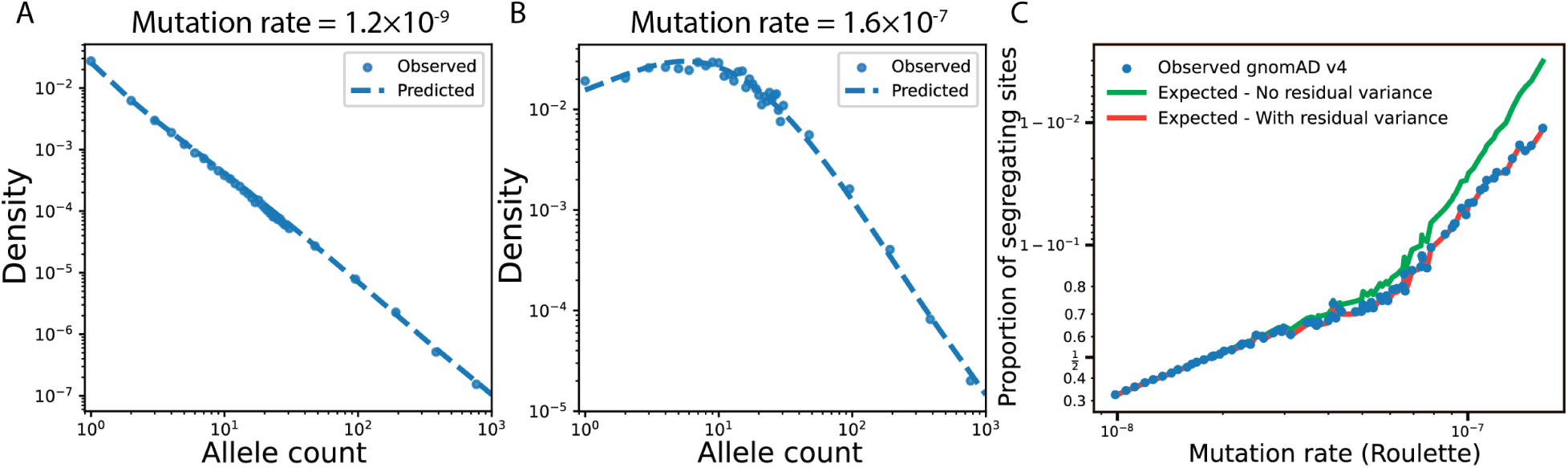
Modeling mutation rates. **A-B.** We show the observed SFS for synonymous variants in the Non-Finnish European (NFE) population from gnomAD v4, stratified by Roulette mutation-rate bins corresponding to low (0.02; A) and high (3.219; B) rates. For each bin, we fit the SFS (blue dashed line) using the analytical model under the NFE demographic history [14] and a Gamma distribution of mutation rates (see Methods). The inferred mean mutation rates for these sites are 1.2 × 10*^−^*^9^ (A) and 1.6 × 10*^−^*^7^ (B). The shape of the SFS, which differs between the low and high mutation rates due to recurrence, provides information about the mutation rate. **C.** Although Roulette performs better than other models in determining per site mutation rates, it still has considerable unexplained variance within a mutation rate bin. We show that modeling the uncertainty (orange line) as opposed to excluding it (green line), improves the estimation of probability of segregation, an important statistic used in constraint metrics.

Given demographic history and mutation rate parameters estimated exome-wide, PIES uses a Bayesian framework to infer selection coefficients at the gene and variant levels, and detect genes with enhanced LoF mutation rates.

### Selection against loss-of-function variants

Existing methods to estimate gene-specific selection coefficients for heterozygous LoF variants (*s*_het_) do not readily scale to constantly increasing sequencing datasets. Methods that avoid expensive calculations rely on the approximation that neglects drift [15, 17], which breaks down in large datasets where drift-induced variance in mutation counts approaches sampling variance [15, 48]. Other methods rely on expensive simulations [8, 16] or numerical matrix multiplication [18, 49]. PIES-LoF offers a computationally efficient scalable approach (Supplementary Figure S6 and Supplementary Methods).

Even in large datasets, LoF mutation data for short genes are sparse, motivating a Bayesian design. Similarly to earlier developed GeneBayes, PIES relies on prior distributions of selection coefficients for every gene. It parametrizes this distribution as a log-normal with gene-specific means adopted from GeneBayes and the variance being a global parameter shared across genes. We found that using a single global variance leads to stable results across benchmarks while reducing the possibility of overfitting the prior (see Supplementary Information for details). To demonstrate the effect of selection, we plot the SFS for two genes (*SYNE2* and *ABCA10*) that experience different levels of negative selection against LoF variants (Figure 3A). *SYNE2*, associated with muscular dystrophy [50], exhibits strong negative selection while *ABCA10* is tolerant to LoF mutations [51]. Higher selective constraint is characterized by a faster exponential fall-off in SFS. PIES-LoF provides posterior distributions of *s*_het_ for 17,052 autosomal genes. In our benchmarks, we use 90% lower bound estimates (Supplementary Table S3).

**Figure 3:**
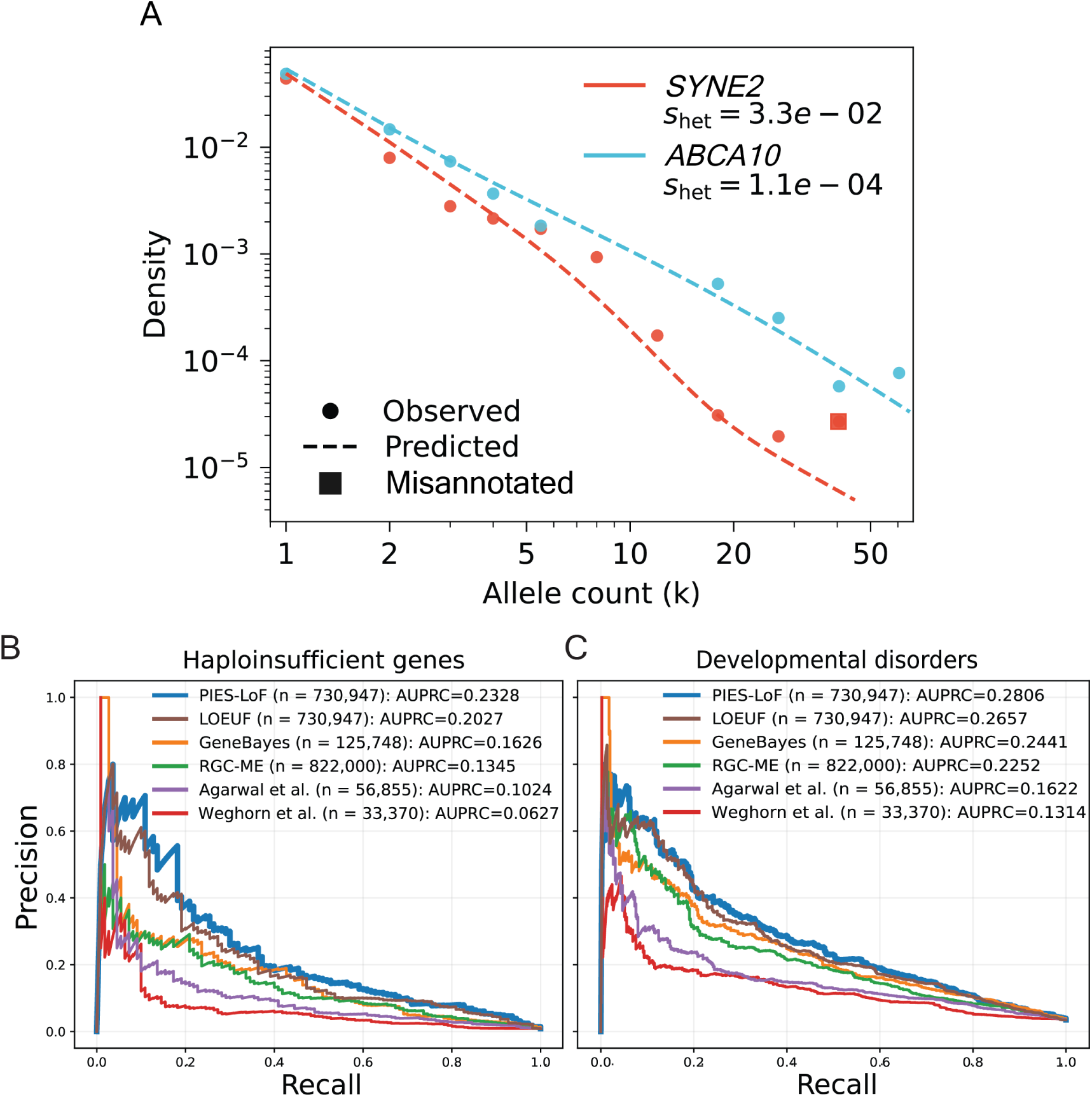
Selection against LoF variants. **A.** We show the observed SFS for LoF variants in the NFE population from gnomAD v4, for two genes - *SYNE2* (red) and *ABCA10* (blue). For each gene, we show PIES-LoF model fits to the SFS (dashed line) obtained using the NFE demographic history [14] and calibrated mutation rate distribution (see Methods). The PIES-LoF framework accounts for potential misannotation of LoF variants (see Methods and Supplementary Table S4). Variants in *SYNE2* with misannotation probability greater than 0.999 are highlighted. **B-C.** We evaluate PIES-LoF against existing s_het_ estimates - [8, 16–18], and the heuristic constraint method - LOEUF applied to gnomAD v4 [5]- in their ability to pick haploinsufficient genes (B) and genes involved in developmental disorders (C). We use the area-under the precision-recall curve (AUPRC) statistic to benchmark the methods (see Data analysis in Supplementary Information).

In order to access the improvement of the s_het_ estimates due to the larger sample sizes in gnomAD v4, we compare PIES-LoF with previous s_het_ estimates [8, 16–18]. We benchmark the estimates in their ability to identify strong selection and disease genes such as haploinsufficient genes [52] and genes involved in autosomal dominant monogenic developmental disorders [53]. We found that the improvement over GeneBayes, computed on gnomAD v2, based on the area-under the precision-recall curve (AUPRC) is significant (difference in AUPRC, p-value = 0.0078, Figures 3B, C; see Data analysis for details). Supplementary Figures S3A and S3B also show the sample size independent comparison of PIES-LoF with GeneBayes on the smaller gnomAD v2 sample sizes. Figures 3A, B also shows the performance of heuristic LOEUF constraint metric computed on gnomAD v4.

These results serve as a proof-of-concept that PIES is scalable to large datasets like gnomAD v4 and can provide reliable estimates.

### PIES identifies genes with enchanced LoF mutation rates

PIES identifies mutation hotspots associated with LoF variants. As noted in the Introduction, one mechanism causing such hotspots is positive selection in spermatogonial cells leading to formation of clones in sperm (Figure 4A). Presence of such germ cell clones leads to an apparent dramatic increase of the human *de novo* mutation rate [24]. LoF variants generated by mutations in these genes will result in variants segregating in the population (even if at low frequencies for genes under purifying selection). The effect of recurrent mutations on SFS (Figure 2B) provides a statistical instrument to identify hotspots of mutation rate from population sequencing data. Recurrent mutations reduce the relative frequency of singletons (variants observed in a single heterozygote) and increase frequency of rare non-singleton variants. For very high mutation rates, the shape of SFS becomes non-monotonic with a clear maximum. The changes in SFS are illustrated in Figure 4A and Supplementary Figure S4. This makes CES potentially detectable from population genomic data alone.

**Figure 4:**
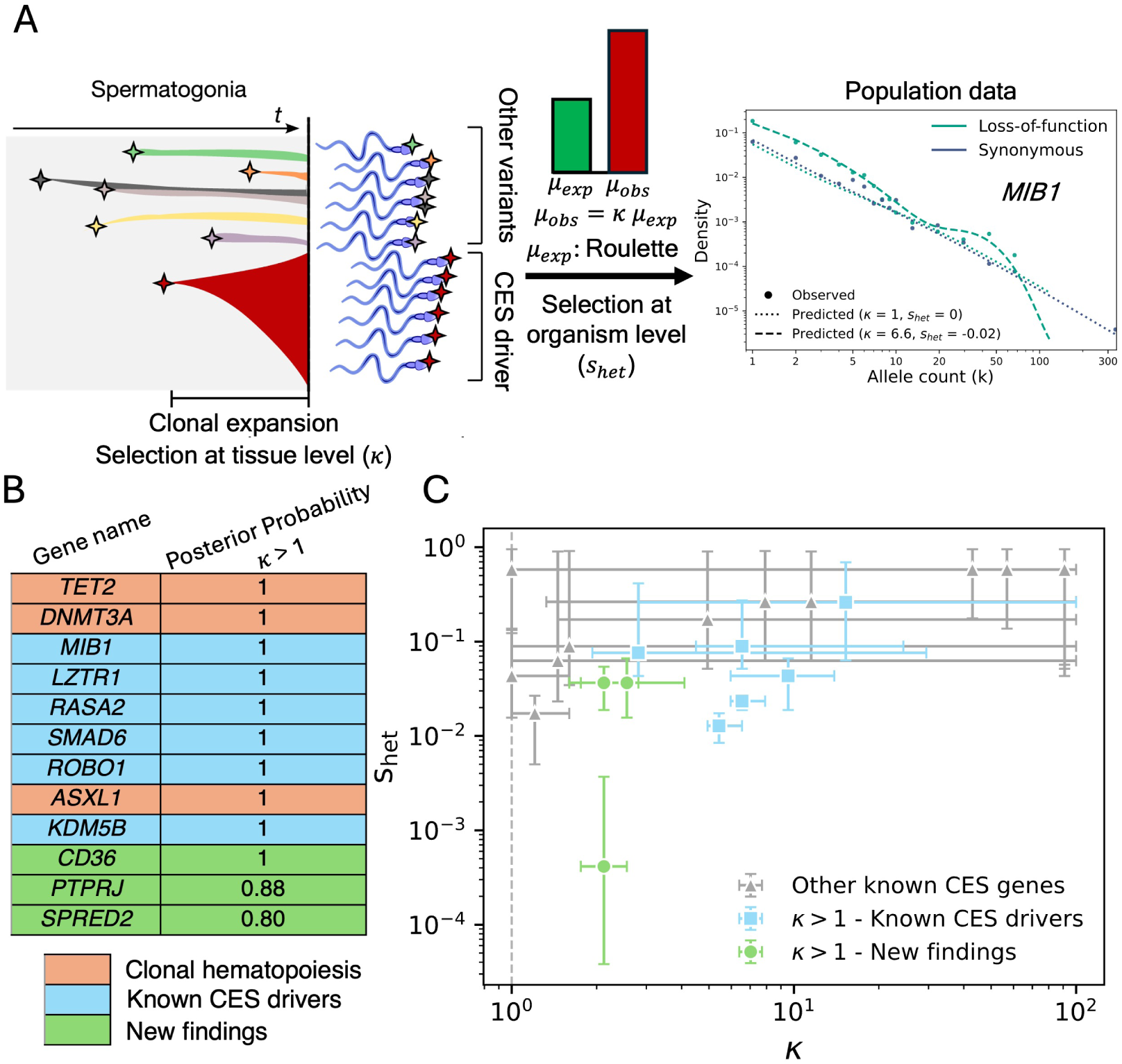
Genes with elevated LoF mutation rates. **A.** Schematic illustrating how clonal expansion in spermatogonia (CES) can lead to an apparent increase in mutation rates for LoFs. We show the SFS of *MIB1*, a known CES driver [24, 25]. **B.** List of genes along with their posterior probability of *κ* to be greater than 1 according to PIES. We divide the gene list into three categories - 1. Genes involved in clonal hematopoiesis - *TET2*, *DNMT3A*, and *ASXL1*; 2. Known CES genes - *MIB1*, *LZTR1*, *RASA2*, *SMAD6*, *ROBO1*, and *KDM5B*; 3.

Apparent mutation rate (*µ_obs_*) in CES driving genes is defined by the “raw” mutation rate (due to DNA damage or replication errors) and the strength of clonal selection in spermatogonia cells. The strength of clonal selection is expected to be identical for all LoF variants in the same gene. Differences in mutation rate among LoF sites in the same gene are only determined by the “raw” rate. This “raw” mutation rate is still expected to be predicted by Roulette (*µ_exp_*). To model clonal selection, we introduce a gene-specific parameter *κ* that is a multiplier of the Roulette predicted mutation rate for LoF mutations. This allows aggregating all LoF variants within a gene to jointly identify mutation hotspots (genes with high *κ*) and the selection coefficient acting against it at the organism level (negative s_het_) (Figure 4A). We use a Bayesian approach to compare the model with LoF mutation rates fixed to the baseline PIES model (*κ* = 1) with a model that allows higher values of *κ* (*κ >* 1). We report Bayes Factor in favor of *κ >* 1 and posterior probability with a prior assuming LoF in 1% of all genes are mutational hotspots (Supplementary Table S5).

The application of PIES to the gnomAD v4 dataset identified 12 genes with an elevated LoF mutation rate, defined by a posterior probability of *κ >* 1 exceeding 0.8 (Figure 4B). Three of them (*TET2, DNMT3A, ASXL1*) corresponded to genes known to drive clonal hematopoiesis and were excluded from consideration. This is because blood is the source of DNA in gnomAD, thus, mutations in clones in blood could be misinterpreted as single nucleotide variants (some of these genes are expressed in spermatogonia and may be genuine drivers of CES) [54]. Six of the remaining genes have been reported earlier from the analysis of *de novo* mutations or direct sperm sequencing [24, 25]. The remaining three genes are either excellent or plausible biological candidates. *SPRED2* is a negative regulator of the MAPK pathway, the major pathway involved in CES [55]. *PTPRJ* is a protein tyrosine phosphatase receptor known to act as a tumor suppressor [56]. *CD36* is a less obvious but still plausible candidate. It is an immune molecule that is involved in lipid uptake that can fuel cell division. It has been shown to play a role in cancer and stem cell metabolism [57]. In the context of spermatogenesis, its role in Sertoli cells, which are responsible for phagocytosis of germ cells, is also of potential interest [58].

Candidate genes with increased LoF mutation rates - *CD36*, *PTPRJ*, and *SPRED2*. **C.** We plot the joint maximum likelihood estimates of *κ* and s_het_ obtained using PIES. Error bars denote 95% confidence intervals derived from profile likelihoods, defined by a log-likelihood drop of 1.92 from the maximum. We plot the known CES drivers that PIES also identifies, new findings, and previously discovered CES drivers [24] not identified by PIES.

We note that some genes previously reported to be involved in CES through the LoF mechanism appear just below our Bayes factor threshold (*TNPO3, GIGYF1, NF1*) [24].

A previous study [24] reported additional 16 genes as potential drivers of CES. Figure 4C shows maximal likelihood estimates of *κ* and s_het_ alongside confidence intervals for these genes and genes identified by PIES. Most of the previously reported genes have high maximal likelihood estimates of *κ* according to PIES, although very wide confidence intervals result in relatively low posterior probabilities. The reason for such wide confidence intervals is the high selection strength (high *s*_het_) limiting the number of segregating variants detectable in gnomAD v4 and, consequently, the confidence of PIES estimates. For three previously identified genes (*TNPO3, GIGYF1, NF1*), confidence intervals do not cross zero but posterior probabilities fall just below our prespecified threshold. Two previously characterized genes (*BCAS3, NARS1*) with low maximum likelihood *κ* estimates cause recessive neurodevelopmental disorders. The PIES model ignores a possibility of recessive selection and may be mis-specified for such genes.

### Selection against missense variants

Estimation of the effect of missense variation is of primary interest because it is the most common type of coding variation and because it is difficult to interpret given that the missense variant effects in the same gene may range from completely neutral to strongly deleterious.

Selection on a missense variant is determined by two factors: (i) the effect of the variant on protein properties such as structure and function, and (ii) the effect of a hypomorphic loss of function (or change of function) of the protein on the organism fitness. Most computational and experimental methods for pathogenicity prediction [33–36, 39–45, 59] focus on the former. However, a small damaging effect in a protein of high importance may be more consequential than a large effect on a less essential gene. We illustrate this point (Figure 5B) by comparing computational predictions (by AlphaMissense) across genes with different LoF selection coefficients in their ability to identify mutations involved in neurodevelopmental disease. Missense mutations that have high values of AlphaMissense score in weakly selected genes show a lower association with neurodevelopmental disease [19] compared to mutations with smaller AlphaMissense values but in more constrained genes.

**Figure 5:**
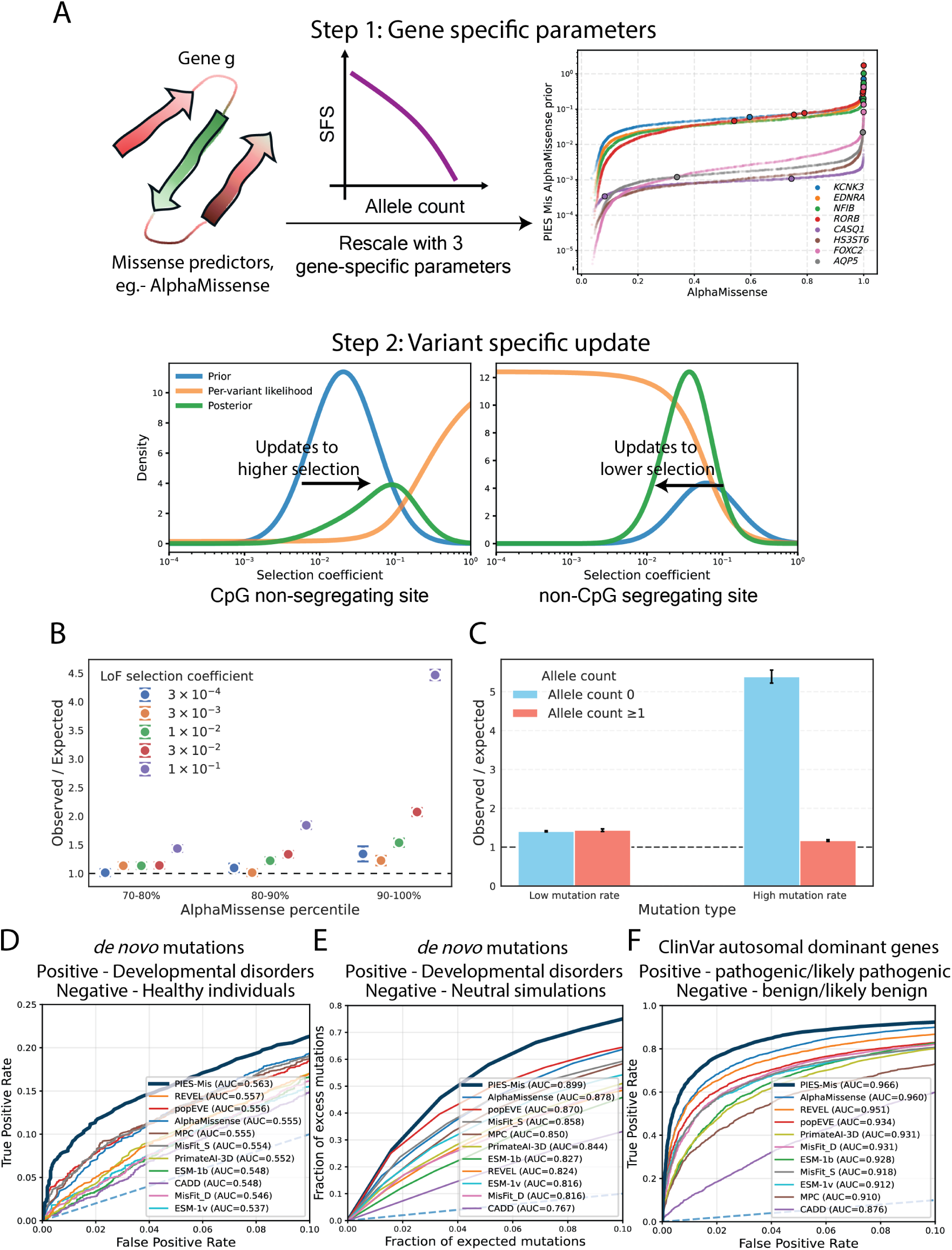
Selection against missense variants. **A.** Schematic of the PIES-Mis framework for estimating selection against missense variants. In step 1 of PIES-Mis, for each gene, we rescale existing missense effect predictors using the observed missense SFS within the gene. We show examples of this calibration based on AlphaMissense for eight genes with similar AlphaMissense distribution. We also label the pathogenic/likely pathogenic labeled ClinVar variants in these genes (circles). We find an increased incidence of pathogenic variants in genes that are upweighted by PIES-Mis method. In step 2, we use the log_10_(s_het_) distribution inferred in step 1 as prior and update it using the variant-specific likelihood to obtain the posterior distribution. Two illustrative examples are shown, left: A non-segregating CpG variant (chr1:1267890:C:T, allele count in gnomAD v4 = 0, mutation rate = 10*^−^*^7^); right: A segregating non-CpG variant (chr1:1338522:C:A, allele count in gnomAD v4 = 2, mutation rate = 3.6 × 10*^−^*^9^). **B.** We stratify the variants by AlphaMissense score; within the top three deciles, we further stratify by gene-level constraint based on PIES-LoF. For each category, we calculate enrichment as the ratio of the observed number of *de novo* variants in probands with neurodevelopmental disorders [19] to the number expected under neutrality from the Roulette mutation model (see Data Analysis). **C.** For variants in the top 50% by AlphaMissense score, we compare enrichment across categories defined by allele count (0 versus ≥ 1) and mutation rate (below or above 5 × 10*^−^*^8^). **D-F.** We compare performance of missense effect predictors and PIES-Mis across three benchmarks. **D.** Receiver operating characteristic (ROC) curve with *de novo* variants in probands with neurodevelopmental disorders as cases and *de novo* variants in healthy individuals as controls [20, 60] (see Data analysis for details). **E.** Comparison using simulated controls derived from the Roulette mutation model. We plot the cumulative difference between observed and expected counts as a function of cumulative expected counts, normalized by their respective totals across all missense variants (see Data analysis for details). **F.** ROC curve for ClinVar variants in dominant genes, where pathogenic or likely pathogenic variants are treated as cases and benign or likely benign variants as controls (see Data Analysis). Curves are truncated to highlight the low false-positive-rate regime, although AUC is computed over the full range.

To estimate selection coefficients for heterozygous missense variants, PIES-Mis models gene-specific distributions of fitness effects conditional on existing missense effect predictions (Figure 5A - Step 1). We assume that log-transformed missense selection coefficients are normally distributed with the mean being a sigmoid function of the prediction score and a constant variance. We fit gene-specific distributions to the observed SFS of missense variants. For each gene, we obtain three parameters corresponding to the inflection point, the steepness of the sigmoid curve, and the variance. At this step, PIES-Mis effectively rescales prediction scores per gene.

We, then, computed per-variant posterior distribution using the baseline PIES model and the inferred gene-specific selection predictions conditioned on variant effect predictors as priors (in an empirical Bayes fashion). PIES-Mis computes posterior distribution for each individual missense variant with the likelihood dependent on allele frequency conditional on mutation rate (Figure 5A - Step 2). The observed allele count of a variant provides informative updates about selection especially in extreme regimes: for example, the absence of variation at highly mutable sites suggests strong purifying selection [61], whereas the presence of variation at sites with low mutation rates indicates weaker constraint (Figure 5A). The practical utility of considering allele frequency in context of mutation rate is shown in Figure 5C contrasting enrichments of *de novo* mutations in probands with neurodevelopmental disease stratified by mutation rate and allele count observed in gnomAD v4.

PIES-Mis reports medians of both prior (PIES-Mis prior) and posterior (PIES-Mis posterior) selection coefficient distributions as estimates for each missense variant (Supplementary Table 6). We advocate for the use of posterior estimates for practical applications. To benchmark performance of PIES-Mis on practically relevant tasks, we focus on the major application areas for predicted effects of missense mutations. Currently, the main application is in assisting decisions in genetic diagnostics [62]. There is a widespread interest in genetic screening, aiming at the early identification of severe monogenic diseases [63, 64]. The problem for diagnostics is in identifying the pathogenic mutation(s) on the background of other variants in the same individual while screening must compare putative pathogenic mutations against others in the population. In both applications, from the statistical perspective, both pathogenic and benign genetic changes are sampled proportional to the mutation influx. We, therefore, performed two benchmarks using *de novo* mutations in trio cohorts of neurodevelopmental disease [19]. Although not all such mutations are necessarily pathogenic, this set is highly enriched in disease-causing missense mutations, given that ∼60% of monogenic neurodevelopmental disorders are caused by a *de novo* mutation [65]. In the first benchmark, we compared *de novo* mutations in probands with *de novo* mutations in unaffected controls [20, 60]. This models the ability to identify pathogenic diagnostic mutations on the background of *de novo* variation in healthy individuals. In the second benchmark, we compared *de novo* mutations in unascertained probands simulated according to Roulette. This benchmark corresponds to the problem of genetic screening that aims to identify individuals with pathogenic mutations on the general background. As seen in Figures 5D, E (and Supplementary Figures S5B-S5C), both prior and posterior PIES-Mis estimates outperform state-of-the-art methods for predicting pathogenic missense mutations. Although the results presented here are based on AlphaMissense scores, improvements over the original predictors are also observed when popEVE and ESM-1b are used to inform priors (Supplementary Figure S5B-S5D). As expected, posterior estimates are slightly superior. To highlight the effect of posterior estimates, we computed enrichment of *de novo* mutations in probands with neurodevelopmental disease across the range of mutation rates. Posterior estimates produce the largest gain in enrichment at high mutation rates (Supplementary Figure S5A).

It is common to benchmark missense predictors on the ClinVar dataset that is an amalgamation of variants reported by clinical diagnostic labs, research studies, expert panels, and literature curation efforts [66]. For this benchmark, we restricted ClinVar to genes causing autosomal dominant phenotypes (PIES estimates selection intensity in heterozygotes). Figure 5F shows that PIES-Mis posterior estimates outperform available computational predictors of missense variant effect. As a dataset, ClinVar does not reflect statistical properties of practical applications. Given that sampling of genes is neither uniform nor defined by the mutation influx, re-weighting prediction scores by gene is not going to show any improvement on the ClinVar benchmark. Still, it can be a useful benchmark for PIES-Mis posterior estimates.

## Discussion

Leveraging the scale of gnomAD v4 [5] and a method based on recent results in population genetics [14], we identified mutation hotspots and estimated strength of negative selection for LoF and missense variants. Our approach relies on SFS that carries more information about population genetics parameters than the density of segregating sites used in several popular constraint methods [11, 38]. Another advantage of relying on SFS is that the density of segregating variants is going to continue increasing with sample size [5]. In the limit, owing to the size of the current human population, every variant compatible with life will be detected. This will render the density of segregating sites an uninformative statistic.

Using the SFS from the population genomic data alone, we are able to detect mutational hotspots for LoF alleles. Given that population sequencing data are available at a greater scale than parent-child trio sequencing datasets or sperm sequencing datasets used in previous studies [24, 25], we identify both previously reported and new signals. Most of the findings can be attributed to CES. However, *CD36* does not have an obvious known role in clonal expansions. It is a key player in the development of male germ cells supporting the idea that most mutation hotspots at functional sites of protein coding genes are linked to sperm cells and their progenitors. *CD36* shows lower mutagenic effects than the identified CES genes, but the mutation increase is highly significant due to a large number of LoF alleles because of the relatively weak purifying selection.

The most important practical application of PIES is in predicting pathogenic missense mutations. PIES-Mis treats the results of existing computational predictors as prior information and updates them using population genetic data at both the gene and variant levels. Our current implementation relies on priors based on machine learning methods such as AlphaMissense, ESM-1b, and popEVE but can incorporate any predictors. This basic philosophy of combining protein language and population genetics models has been previously implemented in popEVE and MisFit [37, 38]. At the same time, PIES-Mis is fundamentally different from popEVE because it relies on SFS rather than density of segregating sites and models recurrent mutations. It is also different from MisFit in the way it combines protein language models with per-variant allele counts.

From the statistical perspective, PIES-Mis is tailored to the problem of genetic diagnostics or genetic screening. It assumes that the data correspond to variants sampled proportionally to their mutation rates. Although we expect PIES-Mis to perform well in other settings such as ClinVar re-annotation (Figure 5F) or gene editing experiment guidance, such applications do not align with the assumed statistical model. Another caveat is that PIES-Mis and PIES-LoF estimate selection coefficient in heterozygotes, meaning that it is not applicable to variants with recessive model of inheritance.

The posterior estimates are strongly dependent on allele frequencies of individual variants. Variant predictors are commonly used in genetic diagnostics as evidence independent of allele frequency (with the latter considered a separate superior source of evidence). The growth of both clinical datasets and reference datasets challenges this approach. For rare variation accessible in gnomAD v4, the coupling between allele frequency and mutation rate renders inference based on a fixed allele frequency threshold misleading (Figure 5C). PIES-Mis calibrates the effect of the gene, the damaging effect of the variant on function, and its allele frequency conditional on mutation rate in a coherent way.

Recent progress in characterizing genealogies including efficient methods for reconstructing ancestral recombination graphs potentially enables a complete characterization of human DNA polymorphism beyond SFS [67–70]. Fundamentally, knowledge of complete genealogies supplement allele frequency data with information on the number of recurrent mutations at the site and their allelic ages. Allelic ages and the number of recurrent mutations conditional on frequency have a limited ability to provide a power gain in the regime of strong negative selection [13, 71–73]. At the same time, the number of recurrent mutations conditional on the overall allele frequency is highly informative about mutation rates. Thus, we are enthusiastic about the promise of genealogy reconstruction for estimation of mutation rates which would likely improve jointly estimated selection coefficients.

Our results support the optimistic outlook for the future of sequencing studies, showing that the growth of population sequencing datasets, paired with parallel development in analysis tools, will continue to increase the resolution of identification of functional and disease-associated variants, aiding biological and clinical analysis.

## Competing Interests

K.J.K. is a member of the scientific advisory board of Nurture Genomics.

## Acknowledgements

We thank Harding Luan and Josh Lichtman for early insightful discussions. We thank Remus Stana, Irene Salinas, Daniel J. Balick, and Ivan Adzhubey for helpful discussions and feedback. K.E.S. and K.J.K. are supported by National Institutes of Health (NIH) grant U24HG011450. V.S. is a UTSouthwestern Medical Foundation endowed scholar. S.R.S. is supported by NIH grants R35GM127131, R01MH101244, and U01HG012009.

## Data Availability

Supplementary tables are uploaded to Zenodo: DOI - 10.5281/zenodo.20587462 [74].

## Notes

https://zenodo.org/records/20587463

