## Supplementary Information for "Inference of elevated mutation rates and variant effects using 700k exomes"

### Supplementary Figures

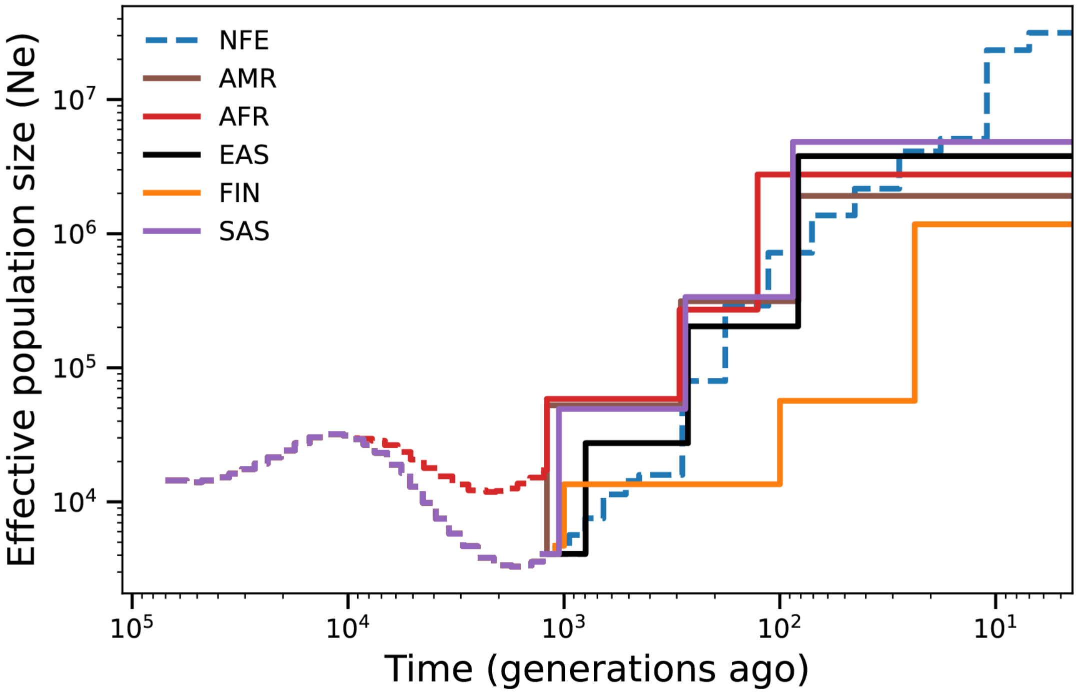


Figure S1: **Demography model used in PIES.** For Non-Finnish European (NFE) ancestry, we use a previously estimated demography model [S1] shown in the figure. For other ancestries in gnomAD v4 - Admixed American (AMR), African/African American (AFR), East Asian (EAS), Finnish (FIN), and South Asian (SAS), we estimate the recent demography using the site frequency spectrum (SFS) of synonymous variants. For these ancestries, we fix the distant past demography (dashed line) [S2]. The recent demography is modeled as piecewise constant with three epochs. We estimate the start times of the epochs and the constant population size in these epochs. The inferred demography for all the ancestries are shown in the figure (solid line). We find rapid population size increase in the recent past for all demographies.

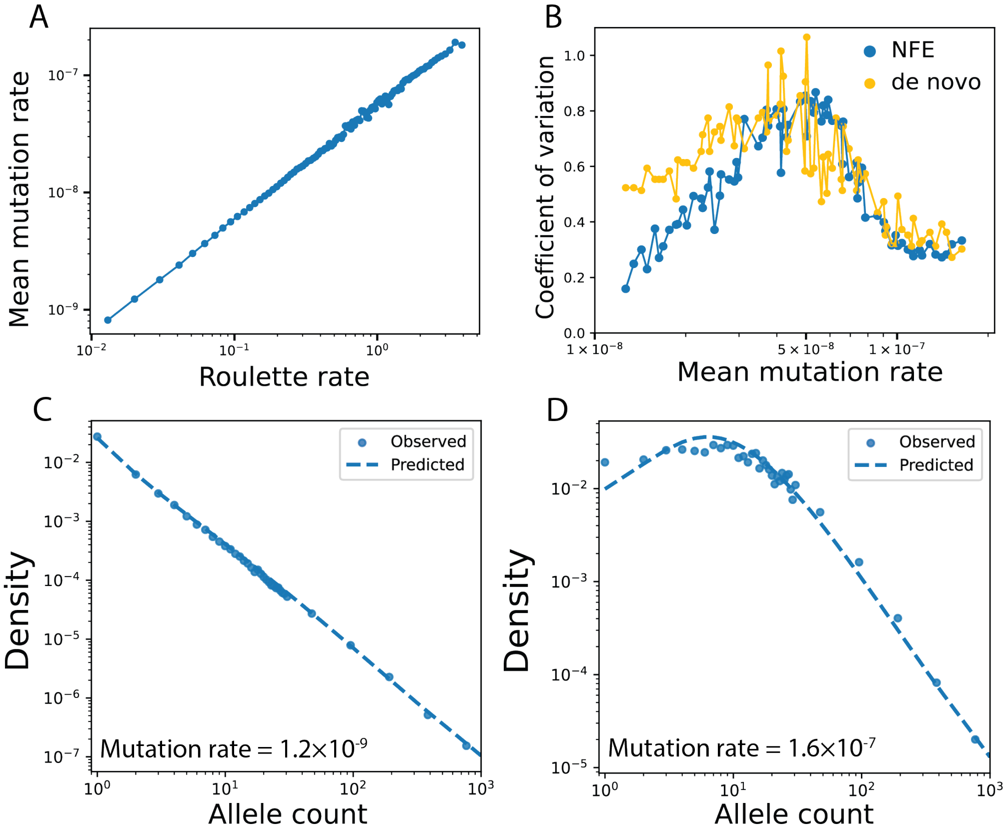


Figure S2: **Modeling mutation rates. A-B.** We estimate mutation rate distributions for each Roulette bin using a three-parameter model within the PIES framework (see Methods). **(A)** The inferred mean mutation rate scales linearly with the corresponding Roulette rate. Thus, PIES effectively rescales Roulette values to per-site, per-generation mutation rates. **(B)** The coefficient of variation (= standard deviation/mean) is shown for different mutation-rate bins. The variance can only be detected for high mutation rate sites. These estimates are consistent with previous values obtained from whole-genome family sequencing data [S3]. **(C–D)** We show the observed SFS for synonymous variants in the NFE population from gnomAD v4, stratified by Roulette bins corresponding to low (0.02; C) and high (3.219; D) mutation rates. A simplified one-parameter mutation-rate model, which does not account for uncertainty, is fit to the observed SFS. While this model provides a reasonable fit at low mutation rates, it fails to capture the SFS at high mutation rate sites. In contrast, the three-parameter model (Figure 2B, main text), which incorporates mutation-rate uncertainty, provides a substantially improved fit in this regime.

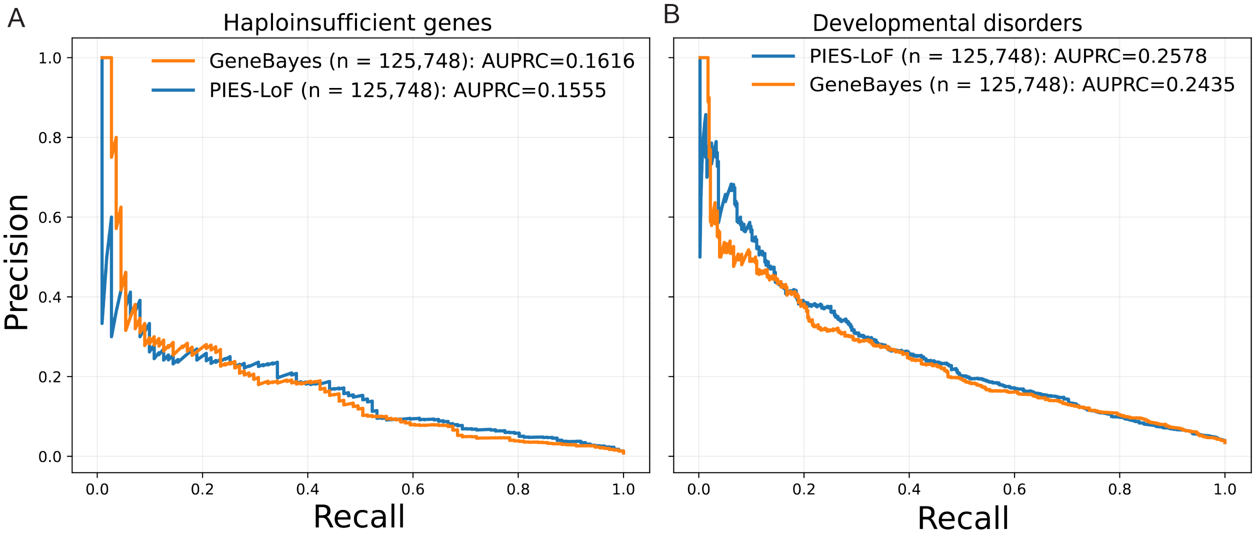


Figure S3: **Benchmarking PIES-LoF applied to gnomAD v2. A-B.** We carry out a sample size independent comparison between PIES-LoF and GeneBayes [S4]. We downsample gnomAD v4 frequencies to gnomAD v2 sample sizes using a hypergeometric distribution and retrained the PIES-LoF model. As in the main text, we compare the two methods in their ability to pick haploinsufficient genes (A) and genes involved in developmental disorders (B). We find the area under the precision-recall curve (AUPRC) to be similar in both datasets (difference in AUPRC, p-value>0.3). Thus, sample sizes play a critical role in improving the loss-of-function (LoF) selection estimates.

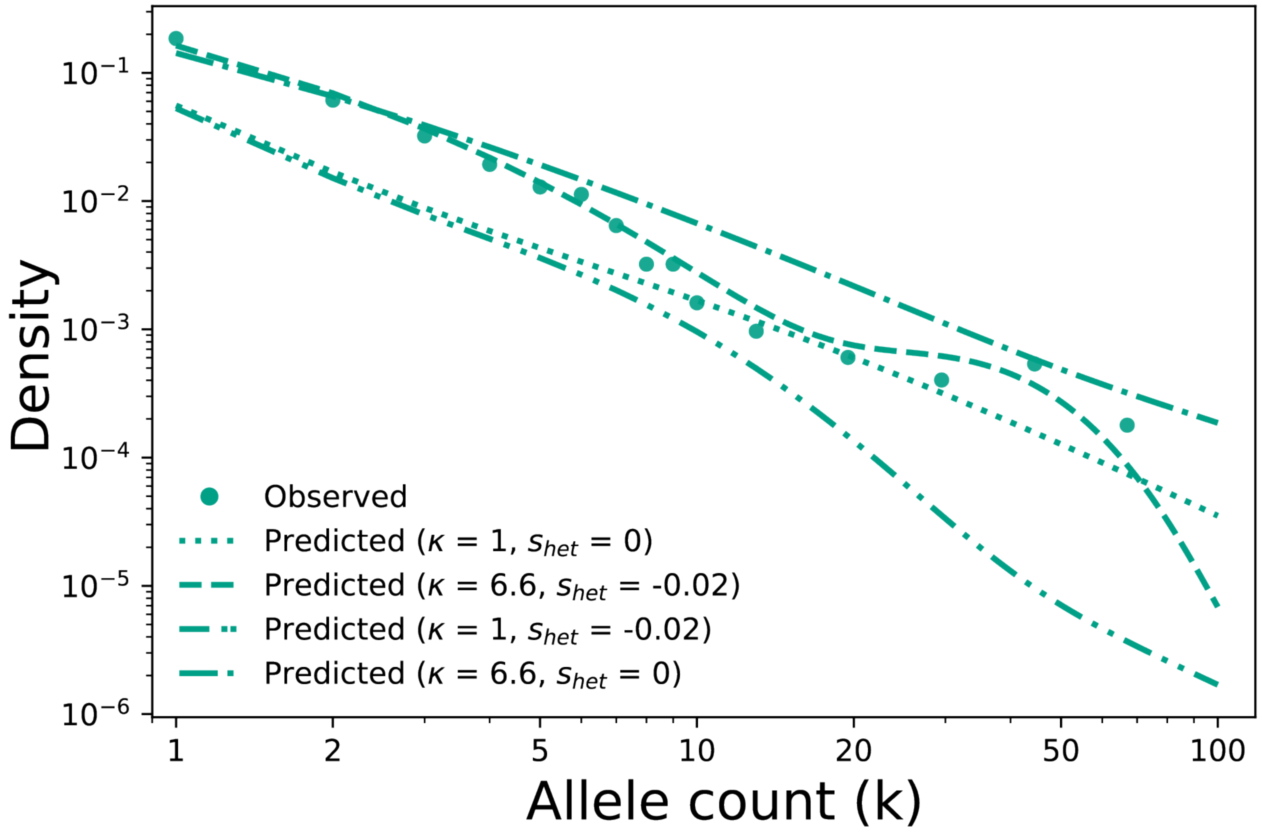


Figure S4: **Model fit to LoF SFS of *MIB1*.** We show the observed SFS of *MIB1* LoFs for NFE ancestry. We also show model fits to the SFS using various values of $\kappa$ (mutation rate inflation factor due to clonal expansion in spermatogonia), and $s_{het}$ (purifying selection against LoFs at organism level). We find the best fit to have $\kappa > 1$, and $s_{het} < 0$. Thus, LoF SFS obtained from population data can be used to detect CES drivers.

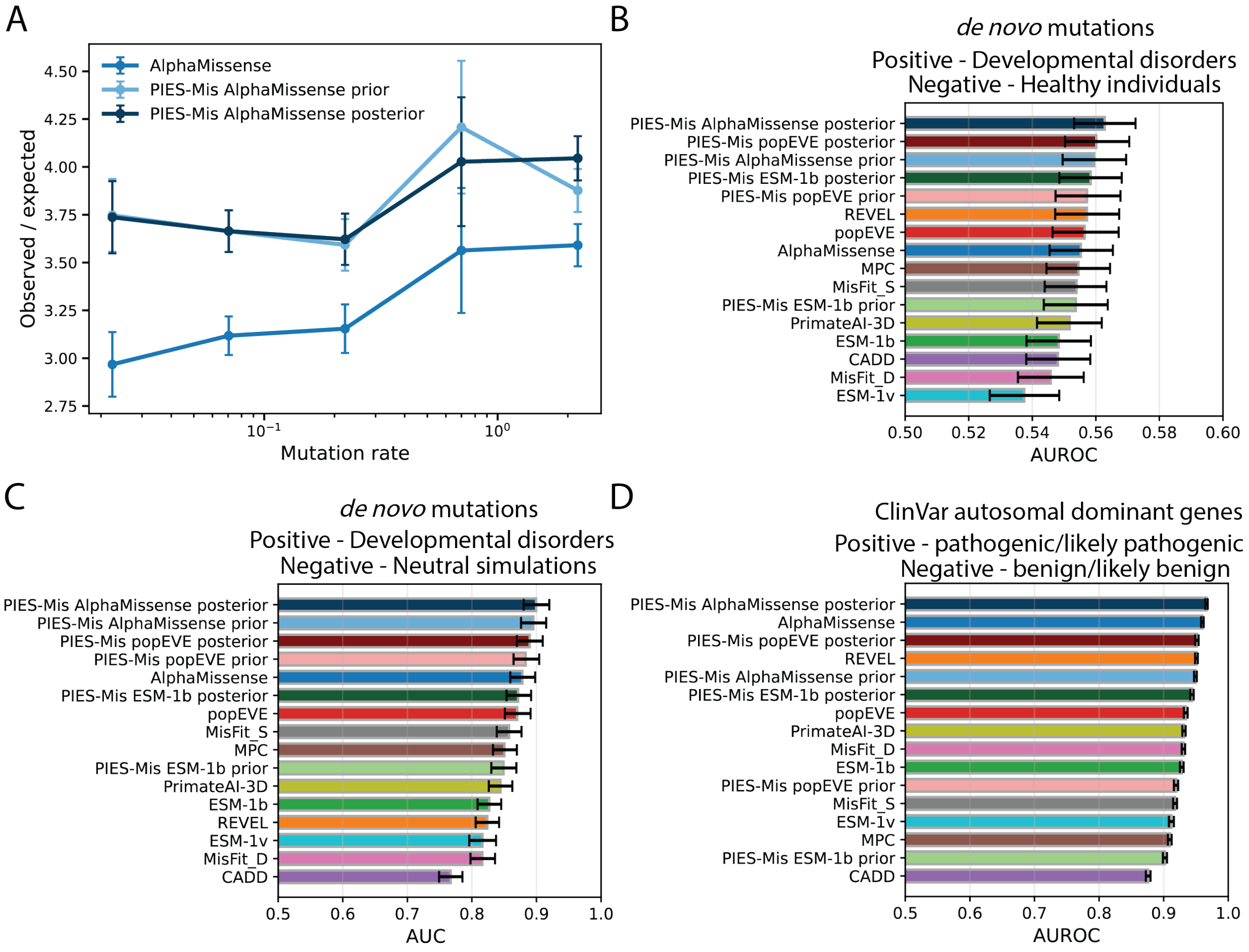


Figure S5: **Benchmarking missense variant effect predictors. A.** We show the improvement from posterior estimates at high mutation rate values. We group the missense variants into five bins based on mutation rates (equal width on a log scale). Within each bin, we compute enrichment - defined as the ratio of observed to expected - among the top 5% pathogenic variants ranked by each predictor. Observed counts correspond to the number of *de novo* variants in probands with neurodevelopmental disorders [S5] while expected counts are derived under a neutral mutation rate model. Assuming Poisson distributed observed counts, the error bars represent $\frac{\sqrt{\text{Observed}}}{\text{Expected}}$. **B-D.** We summarize performance across three benchmarks using the area under the curve (AUC). While full curves for a subset of predictors are shown in the main text, here we report AUC values with 95% confidence intervals obtained via bootstrap resampling. **(B)** Receiver operating characteristic (ROC) AUC for distinguishing *de novo* variants in probands with neurodevelopmental disorders (positive set) [S5] from those in unaffected individuals (negative set) [S6, S7] (see Data Analysis). **(C)** AUC of the cumulative excess metric, defined as the cumulative difference between observed and expected counts as a function of cumulative expected counts, normalized by their totals across all missense variants (see Data Analysis). **(D)** ROC AUC for ClinVar variants in dominant genes, with pathogenic or likely pathogenic variants as the positive set and benign or likely benign variants as the negative set (see Data Analysis).

### Methods

#### Data

Allele counts and allele numbers for each variant are obtained from the publicly available gnomAD v4 dataset [S8]. We use the ancestry labels provided by gnomAD v4. We also use the variant annotations in gnomAD v4, obtained using Ensembl VEP version 105 configured with default parameters for the GRCh38 reference assembly. Variants are classified based on their most severe VEP consequence: "synonymous_variant" for synonymous variants and "missense_variant" for missense variants. A variant is classified as loss-of-function (LoF) if its most severe consequence is one of "splice_donor_variant", "splice_acceptor_variant", or "stop_gained".

We restrict our analysis to autosome genes since Roulette rates are available for only those genes. Additionally, we retain only positions that satisfy the following criteria: (i) "high" and "TFBS" values of the Roulette quality tracks; (ii) positions where allele number $>$ 90% of genome-wide max allele number.

Precomputed raw score tables for each variant effect predictor are first sourced from their corresponding studies [S9–S17] and then standardized to a common representation containing genomic coordinates, alleles, gene identifiers, and predictor-specific scores. The variant effect predictors are merged using genomic coordinates, alleles, and Ensembl gene id. If multiple transcript annotations are available for the same variant, scores are prioritized using a MANE Select first, canonical transcript second, any transcript fallback hierarchy. For predictors where higher scores indicate lower deleteriousness, values are directionally transformed prior to aggregation, ensuring that higher retained scores consistently correspond to greater predicted deleteriousness.

### Population Inferred Estimates of Selection - PIES model

PIES takes rare variation (allele counts $<=5000$ including monomorphic sites) in population data ($\approx$ 1,461,892 haploid exomes in gnomAD v4) as input and estimates parameters encoding changes in population size with time ($N(t)$), mutation rate ($\mu$), and selection coefficient ($s$). It relies on calculating the likelihood of the observed site frequency spectra (SFS) given the population genetic parameters. In the subsequent sections, we will discuss the inference of these parameters sequentially starting with mutation rate and demographic history before moving to gene-specific selection against loss-of-function (LoF) and site-specific selection against missense variants.

The above-mentioned likelihood function of PIES is based on recent theoretical advances to calculate the analytical solution of the sample SFS of rare variants (DR EVIL) [S1]. The sample is obtained by Binomial sampling from the population which evolved under a Wright-Fisher model for given population history, mutation rate and selection intensity. In this rare variant limit, the independent evolution of lineages carrying recurrent mutations at the same site is modeled. Additionally, we assume the dominance coefficient ($h$) to be non-zero; in the rare frequency limit, selection is against heterozygous variants ($s_{het} = s_{hom}h$, where $s_{hom}$ is selection against homozygous variants). We will refer to selection against heterozygous variants as $s$ from hereon. For a given sample of haploid genome of size $n$, we use the probability of observing allele count $k$ given $\mu$, $N$, $s$ parameters ($SFS(k|\mu, s, N)$) as the likelihood in PIES. The analytical solution speeds up the calculation of the predicted SFS compared to the previous state-of-the-art algorithm which relies on matrix multiplication [S18]. Importantly, the computation does not scale with effective population size allowing to estimate the SFS despite large recent population sizes (Supplementary Figure S6). Next, we describe the estimation procedure for different parameters $\mu$, $N$, and $s$ using likelihood functions based on the analytical solution.

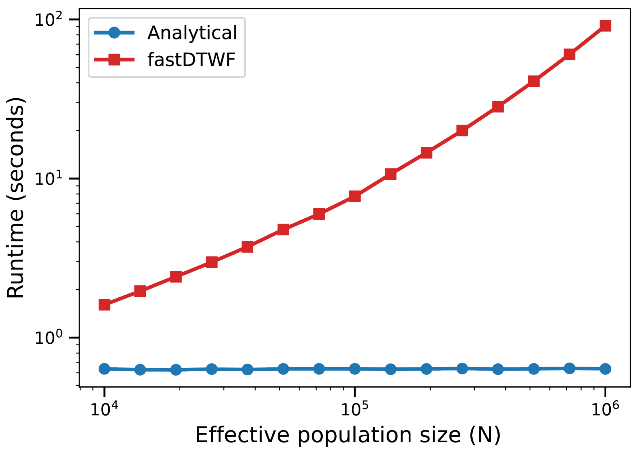


Figure S6: **Runtime of predicted SFS calculations.** We compare the runtime of two approaches for calculating the predicted SFS: (1) fastDTWF, an approximation to the discrete-time Wright–Fisher model [S18], and (2) DR EVIL, which computes the SFS of rare alleles using an analytical solution to the Wright–Fisher model [S1]. We simulate variants with selection coefficient $s_{het} = -10^{-4}$, mutation rate $= 10^{-9}$ per site per generation, and constant effective population size (N). The sample size is set equal to the effective population size. For the analytical method, we limit the maximum allele count at 10,000, independent of N. Each method is simulated three times, and we report the mean runtime. We find that the analytical approach is substantially faster than fastDTWF, with runtime that is effectively independent of N.

#### Calibrating mutation rates

As mentioned in the main text, we supplied PIES with mutation rate predictions from Roulette, which have been shown to provide accurate relative mutation rate estimates [S3]. First, we wanted to scale the relative Roulette rates using population data by fitting the SFS of synonymous sites in each Roulette bin. Thus, we estimate only one parameter - scaled mutation rate $\mu_R$ for each Roulette bin. We assume that the synonymous sites have $s = 0$ and a previously estimated demography $N_{\text{NFE}}$ of Non-Finnish Europeans (NFE) [S1]. The likelihood ($L(\mu_R|k, s, N_{\text{NFE}})$) in this case is,

$$L(\mu_R|k, s = 0, N_{\text{NFE}}) = \prod_{i=1}^{n_{\text{MR}}} \text{SFS}(k_i|\mu_R, s = 0, N_{\text{NFE}}), \tag{S1}$$

where $k_i$ is the allele count of the $i^{th}$ variant and $n_{\text{MR}}$ are the number of variants in each Roulette bin. The likelihood formulation in eq. S1 assumes independence among variants which is appropriate for rare variants which have minimal linkage disequilibrium. We also use this independence assumption in subsequent sections to calculate demography and selection estimates. We find the maximum likelihood estimate of $\mu_R$ using Nelder-Mead optimization algorithm (scipy in Python). We will also use this algorithm in subsequent sections to calculate the maximum likelihood estimates of other population genetics parameters. The one parameter model explains the genetic variation at low mutation rates (Supplementary Figures S2C, S7C) while it fails at higher mutation rates (Supplementary Figures S2D, S7C).

Although Roulette performs better than other models, it still has unexplained variance within a mutation rate bin [S3]. Additionally, ref. [S19] shows that mutation counts in a 1KB region are bimodal with one of the peaks at monomorphic sites (mutational count = 0) possibly due to technical issues such as variant calling. To account for this, we model the absence of mutations as contributions from a low mutation rate site. Thus, we use a three parameter model for the mutation rate distribution in a Roulette bin (Supplementary Figure

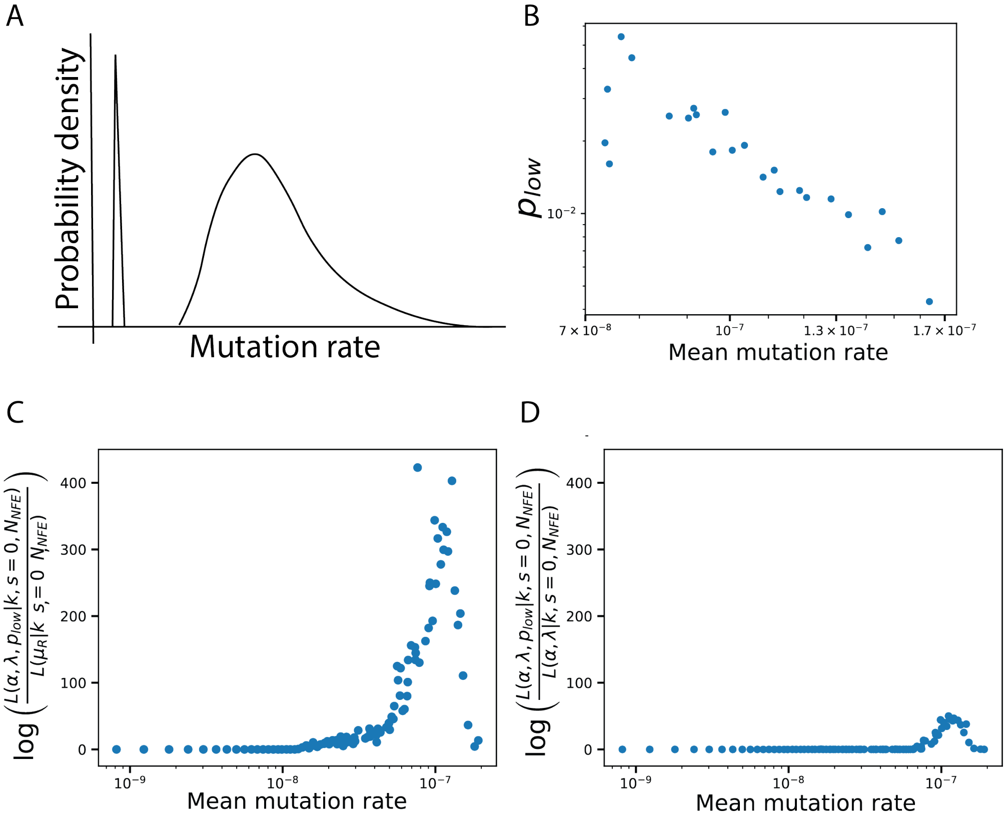


Figure S7: **Mutation rate model comparison.** **A.** Schematic of the three-parameter mutation rate distribution used in PIES for each Roulette bin. The mutation rate is modeled as a mixture of a point mass (with weight $p_{low}$) and a Gamma distribution with shape and rate parameters $\alpha$ and $\lambda$, respectively (eq. S2). **B.** Estimated values of $p_{low}$, which are detectable primarily in high mutation-rate bins. **C-D.** Log-likelihood differences between the three parameter model used in PIES compared to a one parameter model (C) and a two parameter model (D) is plotted for each Roulette bin. Incorporating uncertainty in the mutation rates substantially improves the fit to the synonymous SFS at higher mutation rates (C), with an additional smaller improvement obtained from incorporating $p_{low}$ (D).

S7A),

$$P(\mu) \sim (1 - p_{\text{low}})\Gamma(\mu; \alpha, \lambda) + p_{\text{low}} * \delta(\mu - \mu_{\text{low}}). \quad \text{(S2)}$$

$\Gamma(\mu; \alpha, \lambda)$ is the Gamma distribution with mean $\mu_g = \frac{\alpha}{\lambda}$ and variance $\sigma_g^2 = \frac{\alpha}{\lambda^2}$. $p_{\text{low}}$ is the probability of absence of mutations due to technical issues. $\delta(\mu - \mu_{\text{low}})$ is the Delta function and $\mu_{\text{low}}$ is fixed at $2 \times 10^{-10}$. We use the allele counts of rare (allele counts<=5000) synonymous variants from Non-Finnish Europeans to calculate the likelihood of the three parameters of the mutation rate distribution. The likelihood of the three parameters given the observed allele counts is,

$$L(\alpha, \lambda, p_{\text{low}} \mid k, s=0, N_{\text{NFE}}) = \prod_{i=1}^{n_{\text{MR}}} \left[ (1 - p_{\text{low}}) \int_0^\infty \text{SFS}(k_i \mid \mu, s=0, N_{\text{NFE}})\, \Gamma(\mu \mid \alpha, \lambda)\, d\mu \right.$$
$$\left. +\ p_{\text{low}}\, \text{SFS}(k_i \mid \mu_{\text{low}}, s=0, N_{\text{NFE}}) \right]. \quad \text{(S3)}$$

We use the analytical formula in ref. [S1] to obtain the $\text{SFS}_\Gamma(k_i|\alpha, \lambda, s = 0, N_{\text{NFE}}) = \int_0^\infty \text{SFS}(k_i|\mu, s = 0, N_{\text{NFE}})\Gamma(\mu|\alpha, \lambda)d\mu$. PIES, thus, maximizes the likelihood in eq. S3 to obtain the mutation rate distribution parameters $\alpha$, $\lambda$, and $p_{\text{low}}$. For each Roulette bin, we summarize the maximum likelihood estimates of the three parameter mutation rate model - $4N_0\mu_g$ $(= 4N_0\frac{\alpha}{\lambda})$, $(4N_0)^2\sigma_g^2$ $(= (4N_0)^2\frac{\alpha}{\lambda^2})$, and $p_{\text{low}}$ in Supplementary Table S2. $N_0$ is the initial effective population size and equals 14,448 [S1, S2].

For each Roulette bin, we calculate the mean mutation rate and uncertainty using the estimated values of $\mu_g$, $\sigma_g$, and $p_{\text{low}}$. In line with our expectations, we find a nearly perfect concordance between the mean estimates from PIES and Roulette rates (Supplementary Figure S2A). Residual variance could only be detected for sites with higher mutation rates and it is comparable to previous estimates from *de novo* data [S3] (Figure S2B). We also show the estimates of $p_{\text{low}}$ in high mutation rate regime in Figure S7B where the three parameter model has a higher likelihood compared to the two parameter model that excludes $p_{\text{low}}$

(Supplementary Figure S7D). We will use the estimated mutation rate distribution for each Roulette bin ($\widehat{P(\mu)}$) in the subsequent sections.

#### Estimating recent demography

While Non-Finnish European (NFE) ancestry constitutes the largest ancestry group in gnomAD v4 with a haploid sample size of approximately 1.4 million, there are other ancestries which have sample size greater than 30,000 - Admixed American (AMR, $n = 44724$), African/African American (AFR, $n = 33480$), East Asian (EAS, $n = 39700$), Finnish (FIN, $n = 53420$), and South Asian (SAS, $n = 86258$). The SFS of rare variants provides information about the recent demography history. We use the synonymous allele counts ($s = 0$) from different ancestries to fix the recent demography.

The demographic history is assumed to be a piecewise constant function with the past demography fixed based on a previous study [S2], while we determine the start times, and the effective population sizes of the last three epochs. The non-African ancestries went through an Out-of-Africa bottleneck which we model in the NFE demography used to determine the mutation rate estimates above. We keep the same values for the other non-African demographies until approximately 1200 generations ago (past demography - $N_p(t)$). For AFR ancestry, the population size remained approximately constant in the distant past (also obtained from ref. [S2]). For all of the ancestries, we determine the start time ($\vec{t} = \{t_1, t_2, t_3\}$) and the effective population sizes ($\vec{N} = \{N_1, N_2, N_3\}$) of the non-overlapping last three epochs. Here, $t_1$ represents the start time of the most recent epoch and $N_1$ is the constant effective population size in that epoch while $t_3$ is the oldest epoch that we determine. As mentioned in the main text, we treat the parametric demography model as global rather than gene or variant specific.

In order to calculate $\vec{N}$ and $\vec{t}$, we use $n_D$ low mutation rate synonymous sites as their mutation rate estimates have negligible residual variances and $p_{\text{low}}$ values (see Section "Calibrating mutation rates" in Supplementary text). Hence, we can model the mutation rate as point estimates instead of the three parameter model before. The likelihood function ($L(\vec{N}, \vec{t}|k, s = 0, N_p)$ for estimating the recent demography is,

$$L(\vec{N}, \vec{t}|k, \mu, s = 0, N_p) = \prod_{i=1}^{n_D} \text{SFS}(k_i|\hat{\mu_i}, s = 0, N_p, \vec{N}, \vec{t}), \tag{S4}$$

where $\hat{\mu_i}$ is the estimated population-scaled Roulette value ($\mu_g$) of variant $i$ calculated by maximizing the likelihood in eq. S3. We report the maximum likelihood estimates of $\vec{N}$, and $\vec{t}$ for each ancestry in Supplementary Table S1.

We repeat the procedure described in the previous section to estimate mutation rate parameters for each Roulette bin and ancestry, using the corresponding estimated demographic models. The fitted models provide good agreement with the synonymous SFS across all ancestries, for both low and high mutation-rate bins (Supplementary Figure S8A–E). In addition, we observe a consistent linear relationship between Roulette rates and the inferred mean mutation rates across ancestries (Supplementary Figure S8F).

#### Estimating gene-level selection acting against loss-of-function

Obtaining estimates of the selection coefficient is an important task in human genetics. We will first focus on obtaining gene-level constraint against loss-of-function (LoF) variants. The underlying assumption is that LoFs variants trigger nonsense-mediated mRNA decay (NMD), thus, the effects of all LoFs within the gene are the same. We only consider stop-gained, splice-donor, and splice-acceptor sites as LoF. We pool together all the LoF annotated variants within the gene and calculate the gene-level selection coefficient. Next, we describe the estimation procedure for the LoF selection estimates (PIES-LoF).

For each ancestry, we compute the expected SFS across a grid of selection coefficients and mutation rates. We use ancestry-specific mutation-rate distributions $\widehat{P(\mu)}$ (Equation

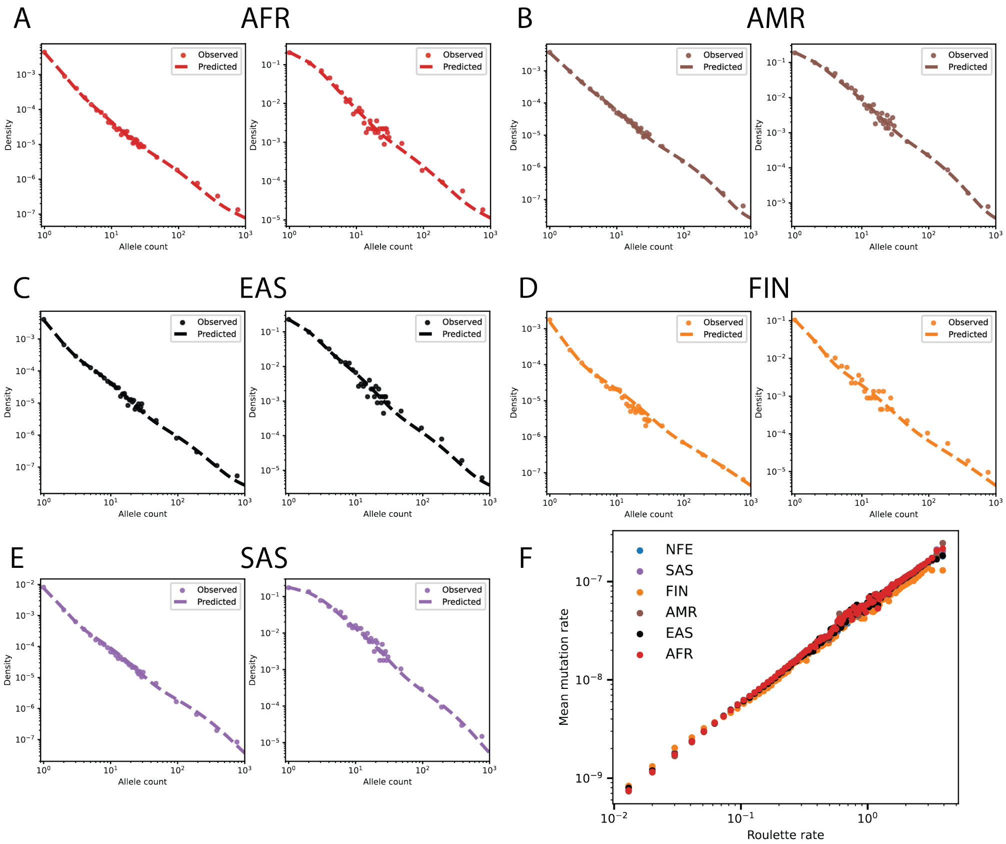


Figure S8: **Fits to synonymous SFS for different ancestries. A-E.** We show the observed SFS for synonymous variants in gnomAD v4 across five ancestries, stratified by Roulette mutation-rate bins corresponding to low (0.02; left) and high (3.219; right) mutation rates. Dashed lines show model fits obtained using PIES, with inferred ancestry-specific demographic histories and a Gamma distribution of mutation rates. The ancestries analyzed are **A.** African/African American (AFR, n = 33480), **B.** Admixed American (AMR, $n$ = 44724), **C.** East Asian (EAS, $n$ = 39700), **D.** Finnish (FIN, $n$ = 53420), and **E.** South Asian (SAS, $n$ = 86258). **F.** Linear scaling between Roulette rates and the inferred mean mutation rates across ancestries demonstrates that Roulette captures relative mutation rates for different populations.

S2) for each Roulette value, together with demographic histories inferred for each ancestry ($\hat{N}$) as described in the previous section. Thus, the resultant $\mathrm{SFS_f}(k|\widehat{P(\mu)}, s, \hat{N})$ is,

$$\begin{aligned}\mathrm{SFS_f}(k \mid \widehat{P(\mu)}, s, \hat{N}) = (1 - \hat{p}_{\mathrm{low}}) \int_0^{\infty} \mathrm{SFS}(k \mid \mu', s, \hat{N})\, \Gamma(\mu' \mid \hat{\alpha}, \hat{\lambda})\, d\mu' \\ +\ \hat{p}_{\mathrm{low}}\, \mathrm{SFS}(k \mid \mu_{\mathrm{low}}, s, \hat{N}),\end{aligned} \tag{S5}$$

where $k$ is the allele count, $\widehat{P(\mu)}$ is the estimated mutation rate distribution corresponding to the Roulette value $\mu$ (eq. S2) and $s$ is the selection coefficient. We remove genes which are involved in clonal expansion of spermatogonia (CES) and clonal hematopoiesis (CH) [S20–S22], as context-based mutation rate models such as Roulette do not account for positive selection in tissues. We restrict the analysis to LoF variants with allele counts $<=$ 5000. In practice, we calculate the SFS across 99 mutation-rate bins defined by Roulette values and a grid of 500 selection coefficients spaced uniformly on a base-10 logarithmic scale between $10^{-6}$ to $10^{2}$ (denoted as set $S$). Note that selection coefficient is on an arbitrary scale and not bounded between 0 and 1, providing relative selection estimates. Higher values of $s$ correspond to a stronger constraint.

Although variants annotated as LoF are typically assumed to trigger NMD and, thereby, result in gene inactivation, this assumption may not always hold. Annotation errors can lead to misclassification of variants as LoF when their functional impact is attenuated or absent. Such misannotations introduce heterogeneity in the true selective effects of variants classified as LoF and may bias the inference of gene-level constraints. We follow the approach in GeneBayes [S4] to account for misannotations of LoFs. We assume that each LoF variant has a certain probability $\mathrm{p}_{\mathrm{mis},f}$ to be misannotated and that the misannotated variants are selectively neutral. The probability value is a global parameter but depends on the functional annotation $f$ of the variant where $f$ can be stop-gained, splice-donor, or splice-acceptor. Hence, the likelihood of selection coefficient acting against the LoF variants ($s_g$)

within gene $g$ for ancestry $a$, and the probability of misannotation $\mathrm{p}_{\mathrm{mis},f}$ given allele counts $k$ are observed is,

$$
\begin{aligned}
L_{a,g}(s_g, \vec{\mathrm{p}}_{\mathrm{mis}} \mid k, \widehat{P(\mu)}, \hat{N}_a, f) = \prod_{i=1}^{n_g} \Bigg[ & (1 - \mathrm{p}_{\mathrm{mis},f_i}) \mathrm{SFS_f}(k_i \mid \widehat{P(\mu_i)}, s_g, \hat{N}_a) \\
& + \ \mathrm{p}_{\mathrm{mis},f_i}\, \mathrm{SFS_f}(k_i \mid \widehat{P(\mu_i)}, s = 0, \hat{N}_a) \Bigg].
\end{aligned} \tag{S6}
$$

The likelihood is calculated using allele counts from a specific ancestry $a$ and for specific gene $g$. Gene $g$ has $n_g$ LoF variants. Each variant $i$ is assumed to be independent. $f$ denotes the functional annotations i.e., stop-gained, splice-donor, and splice-acceptor, for variants in gene $g$ and $\vec{\mathrm{p}}_{\mathrm{mis}}$ is the vector of probability of misannotation consisting of three values corresponding to the values of $f$.

We estimate $\vec{\mathrm{p}}_{\mathrm{mis}}$ globally across genes by profile likelihood method,

$$
\hat{\vec{\mathrm{p}}}_{\mathrm{mis}} = \arg\max_{\vec{\mathrm{p}}_{\mathrm{mis}}} \sum_{g=1}^{G} \max_{s_g \in S} \log(L_{a,g}(s_g, \vec{\mathrm{p}}_{\mathrm{mis}} \mid k, \widehat{P(\mu)}, \hat{N}_a, f)). \tag{S7}
$$

This step treats gene-specific selection coefficients ($s_g$) as nuisance parameters. For fixed values of $\vec{\mathrm{p}}_{\mathrm{mis}}$, we find the maximum log-likelihood values for each gene $g$. So, the profile likelihood function of $\vec{\mathrm{p}}_{\mathrm{mis}}$ equals the sum of maximum log-likelihood values over all genes. The estimated $\hat{\vec{\mathrm{p}}}_{\mathrm{mis}}$ values are those that maximize the profile likelihood function.

We fix $\hat{\vec{\mathrm{p}}}_{\mathrm{mis}}$ values in eq. S6 to obtain the ancestry and gene specific likelihood $L_{a,g}(s_g \mid k, \widehat{P(\mu)}, \hat{N}_a, f, \hat{\vec{\mathrm{p}}}_{\mathrm{mis}})$ as a function of $s_g$. The likelihood $L_g(s_g \mid k, \widehat{P(\mu)}, \hat{N}_a, f, \hat{\vec{\mathrm{p}}}_{\mathrm{mis}})$ for each gene $g$ is obtained by,

$$
L_g(s_g \mid k, \widehat{P(\mu)}, \hat{N}, f, \hat{\vec{\mathrm{p}}}_{\mathrm{mis}}) \approx \prod_{a} L_{a,g}(s_g \mid k, \widehat{P(\mu)}, \hat{N}_a, f, \hat{\vec{\mathrm{p}}}_{\mathrm{mis}}). \tag{S8}
$$

Eq. S8 assumes that allele counts observed in different ancestries are independent. This approximation is justified for rare variants, which are typically of recent origin and therefore unlikely to be shared across populations.

Given a prior distribution $P(s_g)$, we obtain the posterior distribution $P(s_g|k, \widehat{P(\mu)}, \hat{N}, f, \hat{\vec{\mathrm{p}}}_{\mathrm{mis}}$ using the gene-specific likelihood above. We assume a normal prior on $\log_{10}(s_g)$, with mean equal to the gene-specific prior mean reported by GeneBayes [S4] and a fixed standard deviation of 0.5. For each gene, we report the posterior mean and the bounds of the central 95% credible interval in Supplementary Table S3. We additionally report for each variant $i$ in gene $g$, the probability of misannotation ($\mathrm{P}_{\mathrm{mis},i}$) in Supplementary Table S4, calculated as follows,

$$\mathrm{P}_{\mathrm{mis},i}(p \mid \widehat{P(\mu_i)}, k_i, N_{\mathrm{NFE}}, f_i, \hat{\vec{\mathrm{p}}}_{\mathrm{mis}})$$
$$= \frac{\hat{\mathrm{p}}_{\mathrm{mis},f_i}\, \mathrm{SFS_f}\Big(k_i \mid \widehat{P(\mu_i)}, s = 0, N_{\mathrm{NFE}}\Big)}{\hat{\mathrm{p}}_{\mathrm{mis},f_i}\, \mathrm{SFS_f}\Big(k_i \mid \widehat{P(\mu_i)}, s = 0, N_{\mathrm{NFE}}\Big) + (1 - \hat{\mathrm{p}}_{\mathrm{mis},f_i})\, \mathrm{SFS_f}\Big(k_i \mid \widehat{P(\mu_i)}, s = s_{g,\max}, N_{\mathrm{NFE}}\Big)}. \tag{S9}$$

Allele counts are taken from the NFE ancestry, and selection $s_{g,\max}$ is the maximum likelihood selection estimate for gene $g$.

#### Identifying genes with increased LoF mutation rates

Recent studies have found an increased mutation rate of LoFs compared to baseline mutation rate provided by Roulette [S22] due to clonal expansion in spermatogonia (CES). The LoF mutations in these genes provide a selective advantage at the tissue-level resulting in formation of clones. Since these mutations occur in germline cells, they are inherited by offsprings and have signatures in the SFS. We describe the method to obtain genes whose LoFs have higher apparent mutation rate using the population data.

As we restrict to LoFs, we can assume that all LoFs within a gene have the same effect on clonal expansion in tissues as well as purifying selection at the organism level. The increase in mutation rate is due to formation of clones at the tissue level. As mentioned in main text, we model this increase as a factor $\kappa$ over the baseline mutation rate provided by Roulette. For drivers of CES, $\kappa > 1$. We will use the observed LoF SFS of each gene (same as previous section) to jointly infer $\kappa$ and $s$ in this section.

For a given ancestry, we again compute the expected SFS. Unlike the previous section, we have an additional dimension corresponding to $\kappa$ values. We restrict our analysis to NFE ancestry. For each Roulette value and selection coefficient, we obtain the predicted SFS using an equivalent expression as eq. S5,

$$\begin{aligned}\mathrm{SFS}_{\mathrm{f}}(k \mid \widehat{P(\kappa\mu)}, s, N_{\mathrm{NFE}}) = (1-\hat{p}_{\mathrm{low}}) \int_0^\infty \mathrm{SFS}(k \mid \kappa\mu', s, N_{\mathrm{NFE}})\, \Gamma(\kappa\mu' \mid \hat{\alpha}, \frac{\hat{\lambda}}{\kappa})\, d\mu' \\ +\ \hat{p}_{\mathrm{low}}\, \mathrm{SFS}(k \mid \kappa\mu_{\mathrm{low}}, s, N_{\mathrm{NFE}}).\end{aligned} \tag{S10}$$

We restrict the analysis to variants with allele counts $<= 1000$. We exclude single-exon genes based on UCSC Genome Browser annotations, as LoF variants in these genes may escape NMD, leading to heterogeneous effects within a gene. In addition, the grid for selection coefficients is reduced to 100 values, compared to 500 values used in the previous section. The parameter $\kappa$ is evaluated on a grid of 50 values spaced uniformly on a base-10 logarithmic scale between 1 and 100.

Using the SFS in eq. S10, we obtain the likelihood of $\kappa_g$ and $s_g$ for gene $g$ given observed allele counts $k$,

$$\begin{aligned}L_g(\kappa_g, s_g \mid k, \widehat{P(\mu)}, N_{\mathrm{NFE}}, f, \hat{\mathrm{p}}_{\mathrm{mis}}) = \prod_{i=1}^{n_g} \Bigg[ (1-\hat{\mathrm{p}}_{\mathrm{mis},f_i})\mathrm{SFS}_{\mathrm{f}}(k_i \mid \widehat{P(\kappa_g\mu_i)}, s_g, N_{\mathrm{NFE}}) \\ +\ \hat{\mathrm{p}}_{\mathrm{mis},f_i}\, \mathrm{SFS}_{\mathrm{f}}(k_i \mid \widehat{P(\mu_i)}, s=0, N_{\mathrm{NFE}}) \Bigg],\end{aligned} \tag{S11}$$

where, as in the previous section, $n_g$ is the number of LoF variants in gene $g$, $\widehat{P(\mu)}$ is the distribution corresponding to Roulette value $\mu$, $f_i$ is the functional annotation of variant $i$, and $\hat{p_{\mathrm{mis},f_i}}$ is the estimated misannotation probability corresponding to the functional annotations $f_i$. We fix $\hat{\hat{p_{\mathrm{mis}}}}$ to the values estimated in the previous section. Note that the contribution of misannotated variants to the likelihood does not contain $\kappa$ because only true LoF variants drive the clonal expansions.

Next, we use the likelihood function of $\kappa$, and $s$ to compare two models for each gene - M0: $\kappa$ =1, and M1: $\kappa > 1$ i.e., we find the ratio $\frac{P(\mathrm{M1}|\mathrm{Data})}{P(\mathrm{M0}|\mathrm{Data})}$. Using Bayes theorem, we obtain,

$$\frac{P(\mathrm{M1}|\mathrm{Data})}{P(\mathrm{M0}|\mathrm{Data})} = \frac{P(\mathrm{Data}|\mathrm{M1})P(\mathrm{M1})}{P(\mathrm{Data}|\mathrm{M0})P(\mathrm{M0})}. \tag{S12}$$

We set the prior on the models as $\frac{P(\mathrm{M1})}{P(\mathrm{M0})} = \frac{1}{99}$ i.e. 1% of all genes are expected to be CES. We calculate $\frac{P(\mathrm{Data}|\mathrm{M1})}{P(\mathrm{Data}|\mathrm{M0})}$ using the likelihood function $L(\kappa, s|\mathrm{Data}) = L(\kappa, s|k, \widehat{P(\mu)}, N_{\mathrm{NFE}}, f, \hat{\hat{p_{\mathrm{mis}}}})$,

$$\frac{P(\mathrm{Data}|\mathrm{M1})}{P(\mathrm{Data}|\mathrm{M0})} = \frac{\iint P(\kappa, s|\mathrm{M1})L(\kappa, s|\mathrm{Data}, \mathrm{M1})d\kappa.ds}{\int P(s|\mathrm{M0})L(\kappa, s|\mathrm{Data}, \mathrm{M0})ds} = \frac{\iint P(\kappa)P(s)L(\kappa, s|\mathrm{Data})d\kappa.ds}{\int P(s)L(\kappa = 1, s|\mathrm{Data})ds}. \tag{S13}$$

The final equality assumes that $\kappa$ and $s$ are independent, with priors $P(\kappa)$ and $P(s)$,respectively. As in the previous section, we place a normal prior on $\log_{10}(s)$ with mean equal to the gene-specific prior mean reported by GeneBayes [S4] and a fixed standard deviation of 0.5. The prior of $\kappa$ is defined over domain $\kappa > 1$. We choose $P(\kappa)$ such that $\kappa - 1$ follows a Gamma distribution with $P(\kappa) \sim \Gamma(\kappa - 1|mean = 10, variance = 50)$. We note that the set of candidate genes identified (see main text) is robust to reasonable variations in these hyper-parameters. Substituting the functions into eqs. S12 and S13, we compute the posterior probability of model M1 $\left(= \frac{P(\mathrm{M1}|\mathrm{Data})}{P(\mathrm{M1}|\mathrm{Data})+P(\mathrm{M0}|\mathrm{Data})}\right)$ for each gene. We report, for each gene, the posterior probability that $\kappa > 1$ in Supplementary Table S5. Candidate CES genes are defined as those with posterior probability exceeding 0.8. We exclude TEKTIP1, and SBNO2 from the list of novel findings (Figure 4B in main text) as Roulette underpredicts the mutation rates for these genes [S3, S22].

#### Estimating per-variant selection against missense variants

In this section, we describe our approach for estimating selection acting against missense variants. Unlike LoFs, missense variants vary in fitness effects within a gene, thus, we cannot aggregate variants within a gene to obtain a gene-specific selection coefficient.

Existing state-of-the-art computational methods such as ESM-1b and AlphaMissense [S10, S12] infer the functional importance of missense substitutions using properties of the protein such as the protein sequence, evolutionary conservation, and protein structure. Population genomic data is a complementary data modality that can bridge the gap between protein function and missense effects on organism fitness. It motivated us to develop PIES-Mis, which integrates missense effect predictor scores with observed allele counts from gnomAD v4 to infer selection coefficients against heterozygous missense variants.

The computational missense effect predictors capture the relative importance of missense mutations within the protein, however, they may be less useful for comparing gene importance. Therefore, scores for two different genes might be on different scales. To address this, we rescale predictor scores within each gene using the observed SFS of missense variants, enabling consistent interpretation of scores across genes.

First, we assume that the selection for each variant follows a log-normal distribution with mean scaled by missense effect predictors. Given the missense effect predictor score $x$ is,

$$P(\log_{10}(s)|x, c, \beta, \sigma) \sim \text{Normal}\left(z_{\min} + \frac{(z_{\max} - z_{\min})}{1 + e^{-\beta(x-c)}}, \sigma^2\right), \tag{S14}$$

where $c$ is the inflection point, $\beta$ is the steepness of the sigmoid curve, and $\sigma^2$ is the variance of the distribution. $z_{\min}$ and $z_{\max}$ are the bounds of the selection grid (in log-base 10 scale) and are fixed at -6 and 2, respectively. Note that selection $s$ here is scaled by a factor greater than

1. The selection coefficients are evaluated over a finite grid $[10^{z_{\min}}, 10^{z_{\max}}]$. We account for the probability mass of the log-normal distribution that falls outside these bounds. Specifically, the probability mass below $10^{z_{\min}}$ and above $10^{z_{\max}}$ is represented as point masses at the boundaries, with weights given by $F(10^{z_{\min}})$ and $1 - F(10^{z_{\max}})$, respectively, where $F(z)$ denotes the cumulative distribution function of the normal distribution for $z = \log_{10}(s)$.

Eq. S14 simplifies the problem of calculating the selection coefficient for each variant to the inference of three gene-specific parameters. We use the observed SFS of missense variants in gnomAD v4 for each gene to calculate the maximum likelihood estimate of $c$, $\beta$, and $\sigma$. The likelihood $L_g$ of these parameters given observed allele counts $k$ is,

$$\begin{aligned} L_{g,i}(c, \beta, \sigma \mid k, x, P(\mu), N) = &\int_{10^{z_{\min}}}^{10^{z_{\max}}} \mathrm{SFS}_{\mathrm{f}}\Big(k_i \mid \widehat{P(\mu_i)}, s, N\Big) \ \mathrm{LogNormal}(s \mid x_i, c, \beta, \sigma) \ ds \\ &+ \ \mathrm{SFS}_{\mathrm{f}}\Big(k_i \mid \widehat{P(\mu_i)}, 10^{z_{\min}}, N\Big) \ F(10^{z_{\min}}) \\ &+ \ \mathrm{SFS}_{\mathrm{f}}\Big(k_i \mid \widehat{P(\mu_i)}, 10^{z_{\max}}, N\Big) \ (1 - F(10^{z_{\max}}))\,, \end{aligned} \tag{S15}$$

$$L_g(c, \beta, \sigma \mid k, x, P(\mu), N) = \prod_{i=1}^{n_g} L_{g,i}(c, \beta, \sigma). \tag{S16}$$

We consider missense variants that have an allele count$<=$5000. Unlike previous sections, we consider the sum of allele counts from all ancestries as the allele count for a missense variant. We use the NFE demography because NFE samples account for 75% of total samples. We recalculate the parameters for the mutation rate distribution $P(\mu)$ for each Roulette value. The mean mutation rate scales linearly with the Roulette values (Supplementary Figure S9). We use eq. S5 to calculate the expected SFS for different selection coefficients, and 99 mutation rate bins with recalibrated parameters $\alpha$, $\lambda$, and $p_{low}$. The sample size $n$ ($=$ 1461892) is the sum of sample sizes from all ancestries. For each gene, we estimate $c$, $\beta$, and $\sigma$ by minimizing the negative log-likelihood using the Nelder-Mead algorithm implemented in SciPy.

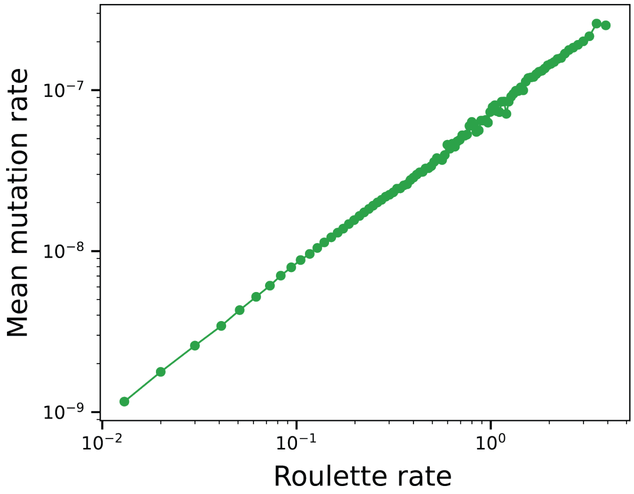


Figure S9: **Mutation rate scaling for combined allele counts.** To estimate selection coefficients for missense variants, we use the SFS constructed from allele counts pooled across all ancestries in gnomAD v4.. We recalibrate the mutation rate distributions for each Roulette bin using NFE demography. As in the ancestry-specific analysis, we observe a linear relationship between Roulette rates and the inferred mean mutation rates. This scaling enables the application of PIES-Mis to SFS derived from combined allele counts.

Although the rescaled predictor scores are comparable across genes, they do not directly leverage allele count information at individual sites. In principle, the observed allele count of a variant can provide informative updates about selection. Thus, we utilize the information of the per variant allele count and the predicted SFS ($\mathrm{SFS_f}$) to obtain the per variant likelihood as a function of selection coefficient. Using the gene-level estimates of $c$, $\beta$, and $\sigma$ as prior (substitute in eq. S14 for each variant $i$ with score $x_i$), we obtain the posterior distribution for each variant $i$,

$$P_{\text{post}}(s \mid k_i, x_i, c_g, \beta_g, \sigma_g) = \begin{cases} \dfrac{\mathrm{SFS_f}(k_i|s)\ \mathrm{LogNormal}\Big(s \mid x_i, \hat{c_g}, \hat{\beta}_g, \hat{\sigma_g}\Big)}{L_{g,i}(\hat{c_g}, \hat{\beta}_g, \hat{\sigma_g})}, & 10^{z_{\min}} < s < 10^{z_{\max}} \\ \dfrac{\mathrm{SFS_f}(k_i \mid 10^{z_{\min}})\ F(10^{z_{\min}})}{L_{g,i}(\hat{c_g}, \hat{\beta}_g, \hat{\sigma_g})}, & s = 10^{z_{\min}} \\ \dfrac{\mathrm{SFS_f}(k_i \mid 10^{z_{\max}})\ (1 - F(10^{z_{\max}}))}{L_{g,i}(\hat{c_g}, \hat{\beta}_g, \hat{\sigma_g})}, & s = 10^{z_{\max}} \end{cases} \tag{S17}$$

where $\hat{c_g}$, $\hat{\beta_g}$, and $\hat{\sigma_g}$ are the gene-specific maximum likelihood estimates.

We report the median estimates from the prior and posterior distribution of each variant $i$. We generate different missense effect prediction tracks for each of the three different predictors - AlphaMissense [S12], ESM-1b [S10], and popEVE [S17]. AlphaMissense scores vary between 0 and 1 and are bimodal. We map AlphaMissense scores within each gene to a normal distribution. For ESM-1b and popEVE, we consider the negative of the scores so that higher values of transformed ESM-1b or popeEVE values denote higher constraints. These per-variant selection estimates are tabulated in Supplementary Table S6.

### Data analysis

#### Precision-recall curves

We evaluate the performance of LoF gene-level constraint estimates using precision–recall (PR) curves on curated benchmark gene sets. We remove known drivers of CES and clonal hematopoiesis from the analysis [S20–S22] as they could appear as false positives in the benchmark sets due to incorrectly estimated mutation rates. The two benchmark sets are the list of haploinsufficient genes [S23] and genes involved in autosomal dominant monogenic developmental disorders [S24]. For each benchmark, genes were ranked based on LoF constraint estimates, and PR curves were constructed by traversing the ranked list from highest to lowest constraint.

Let $U$ denote the set of genes scored by all the methods being evaluated and $B$ be the set of benchmark genes. The performance metrics were computed only with respect to the subset $B \cap U$ (total number = $n_{\mathrm{BU}}$). Thus, genes involved in CES, and CH were excluded. At each rank threshold $r$, true positives (TP) were defined as the cumulative number of benchmark genes encountered up to rank $r$, and false positives (FP) as the number of non-benchmark genes. Precision and recall were then calculated as,

$$\text{Precision}(r) = \frac{\text{TP}(r)}{\text{TP}(r) + \text{FP}(r)}, \qquad \text{Recall}(r) = \frac{\text{TP}(r)}{n_{\text{BU}}}. \tag{S18}$$

We calculate the area under the precision–recall curve (AUPRC) using trapezoidal numerical integration over recall.

To compare the differences in AUPRC of two methods, we used a non-parametric bootstrap procedure. We generated 10,000 bootstrap replicates by resampling genes with replacement from set $U$. For each bootstrap sample $b$, we obtained the difference in AUPRC of the two methods under consideration.

#### Receiver operating characteristic (ROC) curves

We measure performance of missense predictors using area under the ROC curves (AUROC). We use two different datasets - 1. *de novo* variants from trios sequencing, 2. ClinVar variants. At each rank threshold with ordering obtained based on the missense predictor, we find the number of true positives and false positives to obtain the true positive rate (true positive divided by total number of cases) and false positive rate (false positive divided by total number of controls), respectively. The AUROC is the area under the curve. We generate 5000 boostrap replicates by resampling variants with replacement. We calculate the AUROC for each replicate and plot error bars by taking the 95% confidence interval.

##### *de novo* variants

We calculate AUROC values using *de novo* variants obtained from probands with neurodevelopmental disorders as cases [S5] and *de novo* variants from healthy individuals as controls [S6, S7]. We filter the variants to be in autosomes. We remove the variants which were observed both in cases and controls. We overlap the resultant dataset with missense variants which have at least one of the PIES-Mis scores (trained with AlphaMissense/ESM-1b/popEVE as prior). This resulted in total of 26,196 variants.

##### ClinVar variants

ClinVar variant summary was downloaded on March 2, 2026. We filter the variants to be in autosomes and "Assembly" column to be "GRCh38". We keep variants with review status of one star and above. Pathgenic/likely pathogenic variants are regarded as positive set and benign/likely benign as negative set. We overlap the resultant dataset with missense variants that have at least one of the PIES-Mis scores (trained with AlphaMissense/ESM-1b/popEVE as prior). We filter variants to be in genes that cause autosomal dominant phenotypes. The list of dominant drivers of monogenic disease is obtained from OMIM (the full list of genes with "autosomal dominant" labels in OMIM). OMIM was accessed on Oct 12th 2025. We also selected only those genes that had at least one positive and negative label. This resulted in a total of 36,258 variants.

#### Cumulative enrichment curves as a function

In this section, we construct a area under the curve (AUC) statistic that considers the expected number of *de novo* mutations under neutrality as the negative set. To construct this, we order all missense variants by score $x$ in descending order of constraint. Let us define,

$$O_k = \sum_{i \leq k} Y_i, E_k = \sum_{i \leq k} \mu_i, X_k = O_k - E_k. \tag{S19}$$

$O_k$ is the cumulative sum of the observed *de novo* counts ($Y$) until rank $k$ while $E_k$ is the cumulative sum of expected *de novo* counts if the sites are assumed to be neutral.

Observed *de novo* counts ($Y_i$) are obtained from trios sequencing of probands with developmental disorders [S5]. To obtain expected counts ($\mu_i$), we rescale the relative mutation rates determined by the Roulette model using synonymous variants, which are assumed to evolve neutrally. Specifically, the expected count for variant i is given by

$$\mu_i = MR_i \times \frac{\sum_{j \in \text{syn}} Y_j}{\sum_{j \in \text{syn}} MR_j}, \tag{S20}$$

where the scaling factor is computed using observed and expected counts at synonymous sites. We compute the observed to expected ratios in Figures 5A–B and Supplementary Figure S5A as $\frac{\sum_{i \in C} Y_i}{\sum_{i \in C} \mu_i}$, where $C$ denotes the set of variants within a given category.

The total excess of *de novo* mutations across all missense sites is $X_n = O_n - E_n$, where $n$ is the number of missense variants being considered. We define $C_k$ as the fraction of this excess that is captured by sites of rank $k$ and lower,

$$C_k = \frac{X_k}{X_n}. \tag{S21}$$

Let $c_k$ be the cumulative fraction of the mutational opportunity covered up to rank $k$,

$$c_k = \frac{E_k}{E_n}. \tag{S22}$$

We can compute a single summary as the area under this curve,

$$\text{AUC} = \sum_{k=1}^{n} C_{k-1} \Delta c_k. \tag{S23}$$

Upon rearranging the terms in AUC, we obtain,

$$\text{AUC} = \frac{1}{X_n E_n} \sum_{k=1}^{n} X_{k-1} \mu_k = \frac{1}{X_n E_n} \sum_{k=1}^{n} \sum_{i<k} (Y_i - \mu_i) \mu_k = \sum_{i=1}^{n} \frac{(Y_i - \mu_i)}{X_n} \sum_{j>i} \frac{\mu_j}{E_n}. \tag{S24}$$

$$\text{AUC} = \sum_{i=1}^{n} \underbrace{\frac{Y_i - \mu_i}{X_n}}_{\text{average over excess mutations}} \cdot \underbrace{\frac{\sum_{j>i} \mu_j}{E_n}}_{\text{proportion of mutational mass with lower } S \text{ scores}}. \tag{S25}$$

This shows that the AUC calculated from the cumulative excess curve should correspond approximately to the probability that a true excess (causal) mutation has a score greater than one chosen at random from the distribution of new mutations.

Similar to the AUROC calculations above, we calculate the AUC for 5000 bootstrap replicates and plot the 95% confidence intervals as error bars.

### Comparison of selection estimates

In this section, we compare point estimates of LoF selection obtained from PIES-LoF and GeneBayes [S4]. PIES-LoF achieves higher AUPRC values across both benchmarking datasets, largely driven by the increased sample size available in gnomAD v4 (Figures 3B,C). When PIES-LoF is applied to downsampled data matching gnomAD v2 sample sizes, its performance becomes comparable to that of GeneBayes, which was trained on gnomAD v2 (Supplementary Figures S3A,B).

For these comparisons, we use lower-bound estimates of the posterior distribution from both methods. Although lower-bound estimates tend to yield higher AUPRC values in LoF benchmarks (Supplementary Figures S10A,B), there is no *a priori* justification for preferring them as measures of selection strength.

We, next, compare posterior mean estimates between the two methods (Supplementary Figures S10A,B). GeneBayes posterior means yield systematically lower AUPRC values than the corresponding 95% lower-bound estimates from both GeneBayes and PIES-LoF, whereas PIES-LoF posterior means are generally closer to the lower bounds and do not show a similar reduction in AUPRC.

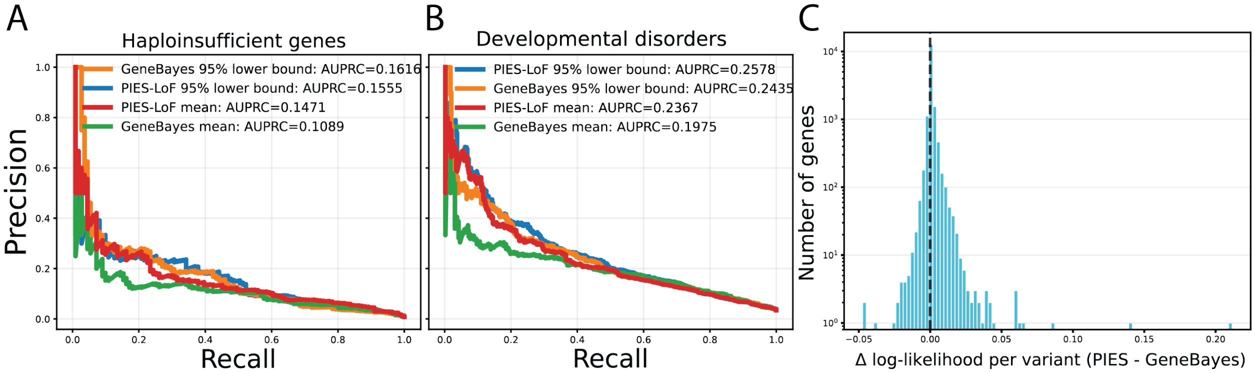


Figure S10: **Comparison between GeneBayes and PIES-LoF. A-B.** We compare mean and 95% lower bound estimates obtained from the posterior distribution of PIES-LoF and GeneBayes. Both methods are applied to gnomAD v2 sample sizes. In benchmarking gene sets of haploinsufficient genes (A) and genes causing developmental disorders (B), we find that AUPRC is significantly lower for the posterior mean from GeneBayes. **C.** Using eq. S8, we compute the likelihood of observing allele counts at posterior means from PIES-LoF and GeneBayes. We plot the histogram of the difference in log-likelihood per variant which is skewed towards positive values.

To further evaluate these estimates, we compare the likelihood of the observed allele counts in gnomAD v2 under each model. For each gene, we compute the likelihood using Equation S8, given mutation rates, demographic parameters, and functional annotations, evaluated at the posterior mean selection estimate. The likelihoods are calculated on data obtained by downsampling gnomAD v4 to match gnomAD v2 sample sizes. We then examine the distribution of the gene-wise log-likelihood differences (PIES-LoF - GeneBayes), normalized by gene length. The resulting distribution is skewed toward positive values, indicating that PIES-LoF posterior means provide a better fit to the observed LoF SFS than those from GeneBayes. We note that PIES-LoF is applied to an independent downsampled version of gnomAD v4, which may introduce a degree of data leakage. Nevertheless, the results in Supplementary Figure S10 suggest that the posterior distribution inferred by PIES-LoF more accurately captures selection effects compared to GeneBayes.
